# Adaptive cellular evolution in the intestinal tracts of hyperdiverse African cichlid fishes

**DOI:** 10.1101/2024.11.28.625862

**Authors:** Antoine Fages, Maëva Luxey, Fabrizia Ronco, Charlotte E.T. Huyghe, P. Navaneeth Krishna Menon, Adrian Indermaur, Walter Salzburger, Patrick Tschopp

## Abstract

Adaptations related to how nutrients are acquired and processed play a central role in the colonization of novel ecological niches and, therefore, in organismal diversification. While the evolution of feeding structures has been studied extensively in this context, the nature of dietary adaptations in the digestive tract remains largely unexplored. Here, we investigate the cellular and molecular basis of dietary adaptations in the massive radiation of cichlid fishes in Lake Tanganyika using comprehensive single-cell transcriptomic data derived from the intestines of 24 endemic cichlid species with distinct habitats and diets. We show that, at the cellular level, dietary adaptations are primarily driven by anterior enterocytes, and that both the relative abundance and gene expression profiles of these cells have evolved in response to rapid dietary specializations. These dietary adaptations are driven by rapidly evolving cell population-specific genes, suggesting that alterations in epithelial cell specification programs and molecular makeup promote ecological diversification.

## Introduction

Adaptations to various types of food are paramount to the survival of animals and key drivers of their evolutionary diversification (Darwin, 1859; Simpson, 1953; Futuyma & Moreno, 1988). These so-called dietary adaptations target all features involved in the localization, uptake, processing, and digestion of food (Karasov, Martínez del Rio & Caviedes-Vidal, 2011), and are manifested in the remarkable diversity of animals in terms of their feeding specializations, foraging behaviors, trophic morphologies, and digestive physiologies (Starck & Wang, 2005; Pough, Janis & Heiser, 2013).

In vertebrates, the nutritional process involves two functionally and anatomically independent units: the feeding apparatus, responsible for the uptake and mastication of food items, and the digestive tract, responsible for enzymatic digestion and nutrient absorption (Pough *et al*., 2013). The importance of adaptive modifications in the feeding apparatus for evolutionary diversification is well documented across animals and particularly evident in cases of adaptive radiations, where new species evolved rapidly from a common ancestor as a consequence of adaptation to distinct ecological niches (Schluter, 2000; Gavrilets & Losos, 2009). Famous examples of adaptations in the trophic apparatus in the ecological context of evolutionary radiations include the varied beaks of Darwin’s finches (Grant, 1986; Abzhanov *et al*., 2006) and the diverse mouth and jaw morphologies of East African cichlid fishes (Fryer & Iles, 1972; Salzburger, 2018; Ronco *et al*., 2021), which, in both cases, match the respective food sources and feeding habits of the species (Schluter, 2000). In contrast, comparatively little is known about adaptations in the intestinal tracts in such outbursts of organismal diversification, except for the association between intestine length, habitat, and diet (Wagner *et al*., 2009). In addition, dietary adaptations during evolutionary radiations have so far mainly been examined at the levels of the gross phenotype (Grant & Grant, 2006; Wagner *et al*., 2009; Ronco *et al*., 2021), genotype (Abzhanov *et al*., 2004; Brawand *et al*., 2014; Lamichhaney *et al*., 2015; El Taher *et al*., 2021), and development (Abzhanov *et al*., 2006; Mallarino *et al*., 2011), but feeding-related adaptations at the cellular level have not been investigated. Yet, the key role that the intestinal tract plays in the exploitation of novel nutritional resources as well as its cellular complexity – comprising multiple cell types involved in digestion, nutrient, and water absorption, hormone production, metabolism, and immunity (Rombout *et al*., 2011; Greenwood-Van Meerveld, Johnson & Grundy, 2017; Elmentaite *et al*., 2021) – suggest that changes in its cellular composition and cell type-specific molecular programs are likely to promote ecological specialization and phenotypic evolution (Luca, Perry & Di Rienzo, 2010; Ocampo *et al*., 2022).

In this study, we scrutinize cellular evolution of the intestinal tract in what is the morphologically, ecologically and behaviorally most diverse adaptive radiation of cichlid fishes, the cichlid fauna of Lake Tanganyika in East Africa. Approximately 250 cichlid species have evolved in this lake from a common ancestor in less than 10 million years (My) (Fryer & Iles, 1972; Rüber & Adams, 2001; Muschick, Indermaur & Salzburger, 2012; Ronco *et al*., 2020, 2021), and have diversified into various trophic niches. The species differ greatly in the morphology of their oral jaws and their pharyngeal jaw apparatus (Ronco *et al*., 2021; El Taher *et al*., 2021; Ronco & Salzburger, 2021) – a functionally and anatomically independent second set of jaws used to mechanically process food (Liem, 1973) – as well as in intestine length and microbial communities (Wagner *et al*., 2009; Baldo *et al*., 2017; Duque-Correa *et al*., 2024). To probe for adaptations at the levels of intestinal tissue composition and cell-type-specific gene expression programs, we capitalize on the power of single-cell RNA sequencing (scRNAseq) data derived from the intestinal tracts of 24 highly ecologically diverse cichlid species from Lake Tanganyika. By examining cellularly resolved transcriptomes in a comparative framework (Tanay & Sebé-Pedrós, 2021; Feregrino & Tschopp, 2022) and in combination with comprehensive eco-morphological data indicative of foraging ecology and dietary niche, we detect molecular signatures of dietary adaptations across multiple layers of biological organization and identify drivers of rapid cell type and tissue evolution in the context of this hyperdiverse vertebrate adaptive radiation.

## Results

### Cellular diversity in the intestines of cichlid fishes

The cellular composition of the intestine is well documented in classical model organisms (Haber *et al*., 2017; Hung *et al*., 2020; Jones *et al*., 2023), but remains unexplored in major ecological and evolutionary study systems such as East African cichlid fishes. We therefore generated scRNAseq data from multiple sections covering the foregut, midgut and hindgut of 24 cichlid species endemic to Lake Tanganyika plus the Nile tilapia (*Oreochromis niloticus)* as an outgroup (Figure 1A). The species were chosen to represent all 13 sub-clades (so-called “tribes”) of the cichlid adaptive radiation in Lake Tanganyika (Ronco *et al*., 2020, 2021), the main ecological niches that the cichlids occupy in this lake as well as the varied dietary specializations that they exhibit, ranging from purely herbivorous/algivorous to omnivorous, to invertivorous, to piscivorous species and even to scale-eating (Muschick *et al*., 2012; Takeyama, 2021; Ronco *et al*., 2021). After mapping the scRNAseq data to the closely related and phylogenetically equidistant Nile tilapia reference genome (Conte *et al*., 2017), data integration, and quality filtering, we obtained 82,261 high-quality cells (mean cells per species: 3304, mean UMI per species: 4109, Figure S1A-D). Based on a combination of marker gene expression information, modules of gene co-expression (Feregrino & Tschopp, 2022), and an *in silico* single-cell-type identifier (Li *et al*., 2020), we defined 24 cell populations in the cichlids’ intestines that belong to nine broad cell-type categories: absorptive epithelial cells (shown in orange in Figure 1), non-absorptive epithelial cells (blue), and cells of the lamina propria, including T cells (light green), B cells (dark green), macrophages (purple), dendritic cells (light purple), neutrophils (dark purple), red blood cells (red), and mesenchymal cells (pink) (Figure 1B-D; Figure S1). The most abundant cell populations detected in the cichlids’ intestinal samples were immune cells, particularly T cells (Figure 1B), consistent with the prominent role that the intestine plays in the immune response of teleost fishes (Rombout *et al*., 2011). Among epithelial cells, defined here by the marker gene *epcam* (Figure 1C), absorptive cells were the most numerous, mirroring patterns reported from mammals (Peterson & Artis, 2014; Haber *et al*., 2017). We identified all known mammalian absorptive and secretory cell types, except for Paneth cells, which have not been observed in other teleosts either (Ng *et al*., 2005; Løkka & Koppang, 2016). We also detected epithelial dendritic cells, so far only reported in the intestinal epithelium of the mouse (Farache *et al*., 2013; Rivera *et al*., 2022) and lysosome-rich enterocytes (LREs), a distinct absorptive cell population involved in protein uptake that has been described in the zebrafish and is absent in post-weaning mammals (Harper *et al*., 2011; Muncan *et al*., 2011; Park *et al*., 2019; Childers *et al*., 2024).

**Figure 1.**
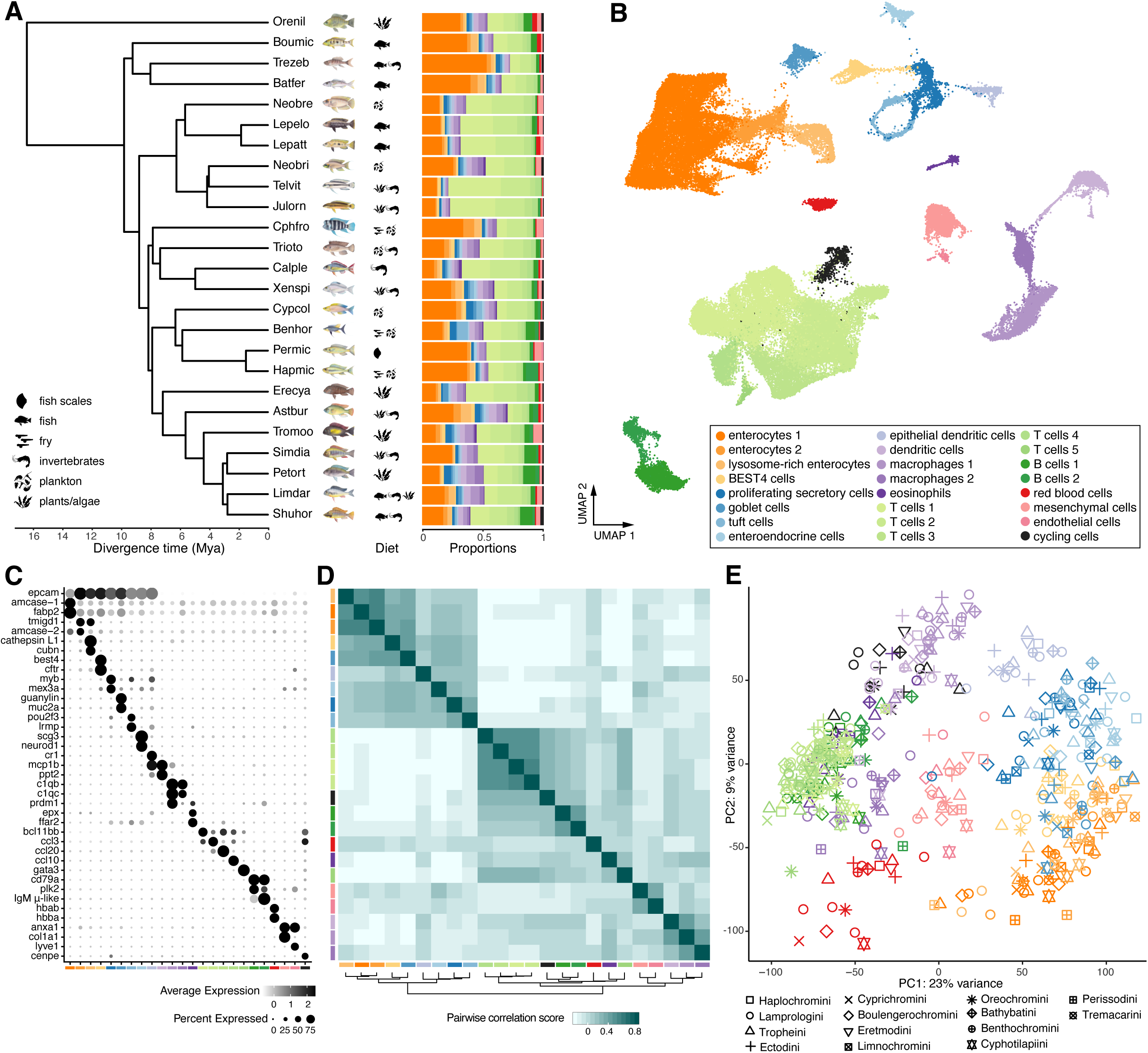
Characterization of the intestines of 25 cichlid fish species at single-cell resolution. (A) Left: Time-calibrated genome-wide species tree of the 25 African cichlid species included in this study, with dietary specializations indicated by symbols. Species names are abbreviated following the nomenclature detailed in Ronco and colleagues (Ronco *et al*., 2021). Right: Relative abundances of cell populations in the intestine samples of each cichlid species. (B) Uniform manifold approximation and projection (UMAP) and cell-population-based clustering of 82,621 cells sampled from the intestines of the 25 cichlid species shown in panel A. (C) Dot plot of selected marker gene expression, used to identify intestinal cell populations. (D) Heatmap of pairwise gene expression correlations between intestinal cell populations. (E) Principal component analysis (PCA) of pseudobulk gene expression variation across all cell populations and species. In all panels, colors refer to cell populations and shapes to the taxonomic clades (tribes) found in Lake Tanganyika. See also Figures S1 and S2 and Table S1.

To approximate the anterior-posterior identity of our epithelial single-cell transcriptomes, we sampled five sections along the intestines of three Lake Tanganyika cichlid species with divergent ecologies and performed bulk RNA sequencing. We then investigated the expression of conserved marker genes specific to different intestinal regions in the mouse and the zebrafish (Figure S2A), and estimated, for each section, the abundance of all cell populations detected in our scRNAseq dataset using a deconvolution approach (Lickwar *et al*., 2017; Chu *et al*., 2022); Figure S2B). We found that the anterior, intermediate, and posterior intestinal regions were enriched in enterocytes 1, enterocytes 2, and LREs, respectively (Figure S2C). Functional characterization of differentially expressed genes through gene ontology (GO) enrichment analysis revealed regional specialization of each of the three enterocyte populations (Figure S2D-G). Thus, although a morphological compartmentalization of their intestines is not apparent in cichlid fishes (Hopperdietzel *et al*., 2014), we show that, just like in other teleost fishes (Calduch-Giner, Sitjà-Bobadilla & Pérez-Sánchez, 2016; Lickwar *et al*., 2017; Park *et al*., 2019), cichlids also feature a molecular and cellular sub-functionalization along the anterior-posterior axis of their intestine.

To further characterize the identified cell populations, we estimated pairwise gene expression correlations between all cell populations on the basis of our scRNAseq data and found that, in general, cell populations cluster by broad cell type. Notably, immune and epithelial cell populations showed the strongest gene expression affinities within their respective groups, indicating a high degree of transcriptional similarity among them (Figure 1D, Figure S1F). We next assessed interspecific gene expression divergence relative to cross-cell-population disparity by calculating pseudobulk values for each gene, cell population, and species. A principal component analysis (PCA) based on the top 10,000 variable genes revealed that each cell-type category and cell population can be well-characterized through the first four principal components (PCs), which collectively explain 41% of the total variance: PC1 (23%) separates epithelial cells from all other cell populations, while PC2 (9%), PC3 (5%), and PC4 (4%) distinguish absorptive from non-absorptive epithelial cells (Figure 1E, Figure S1G,H). The PCA also revealed that cross-species gene expression variation is primarily driven by differences between cell populations and not by differences between species, which is consistent with bulk transcriptome comparisons in various organs across mammals (Berthelot *et al*., 2018) and cichlids (El Taher *et al*., 2021). This suggests that at the within-tissue level, and after approximately 10 million years of divergence, cell population-specific gene expression differences dominate over species-level differences, thus permitting the identification of homologous cell populations and the detection of potentially adaptive species-specific gene expression patterns therein.

### Transcriptome evolution of the cichlids’ intestinal cell populations

To characterize the evolutionary trajectories of the intestinal cell populations detected in cichlid fishes, we first investigated gene expression divergence patterns across species for the 18 cell populations containing at least 20 cells in 17 species. For each of these cell populations, we computed pseudobulk values and calculated mean squared gene expression distances across all species pairs (Figure S3A). We found that the patterns of gene expression divergence in relation to evolutionary distance (calculated as substitutions across the entire genome across species pairs) closely resembles an Ornstein-Uhlenbeck process, with gene expression divergence increasing rapidly at short evolutionary distances, but saturating across longer distances (Hansen, 1997); Figure 2A). This is in line with previous findings in mammals and fruit flies (Bedford & Hartl, 2009; Chen *et al*., 2019) and corroborates the common view that gene expression in animals is constrained by stabilizing selection (Fay & Wittkopp, 2008; El Taher *et al*., 2021; Hill, Vande Zande & Wittkopp, 2021). Interestingly, however, absorptive epithelial cells exhibited greater gene expression differences across species than any other cell population, indicating increased divergence rates in these nutrient-absorbing cells (Figure 2A,B). In contrast, epithelial proliferating cells and non-epithelial cells, in particular immune cells, displayed consistently low inter-specific divergence in gene expression (Figure 2B). This was confirmed by an independent machine-learning framework quantifying the extent of informative species-specific differentiation within each cell population and, therefore, how transcriptionally distinct cell populations are from species to species (Skinnider *et al*., 2021); Figure 2C). The differences in interspecific divergence across cell populations were more marked in protein-coding genes (PCGs) compared to both long non-coding RNAs (lncRNAs) and transcription factors (TFs) (Figure S3B-D), while cross-species divergence was lower and increasing slower with time in PCGs and TFs than in lncRNAs (Figure S3E,F).

**Figure 2.**
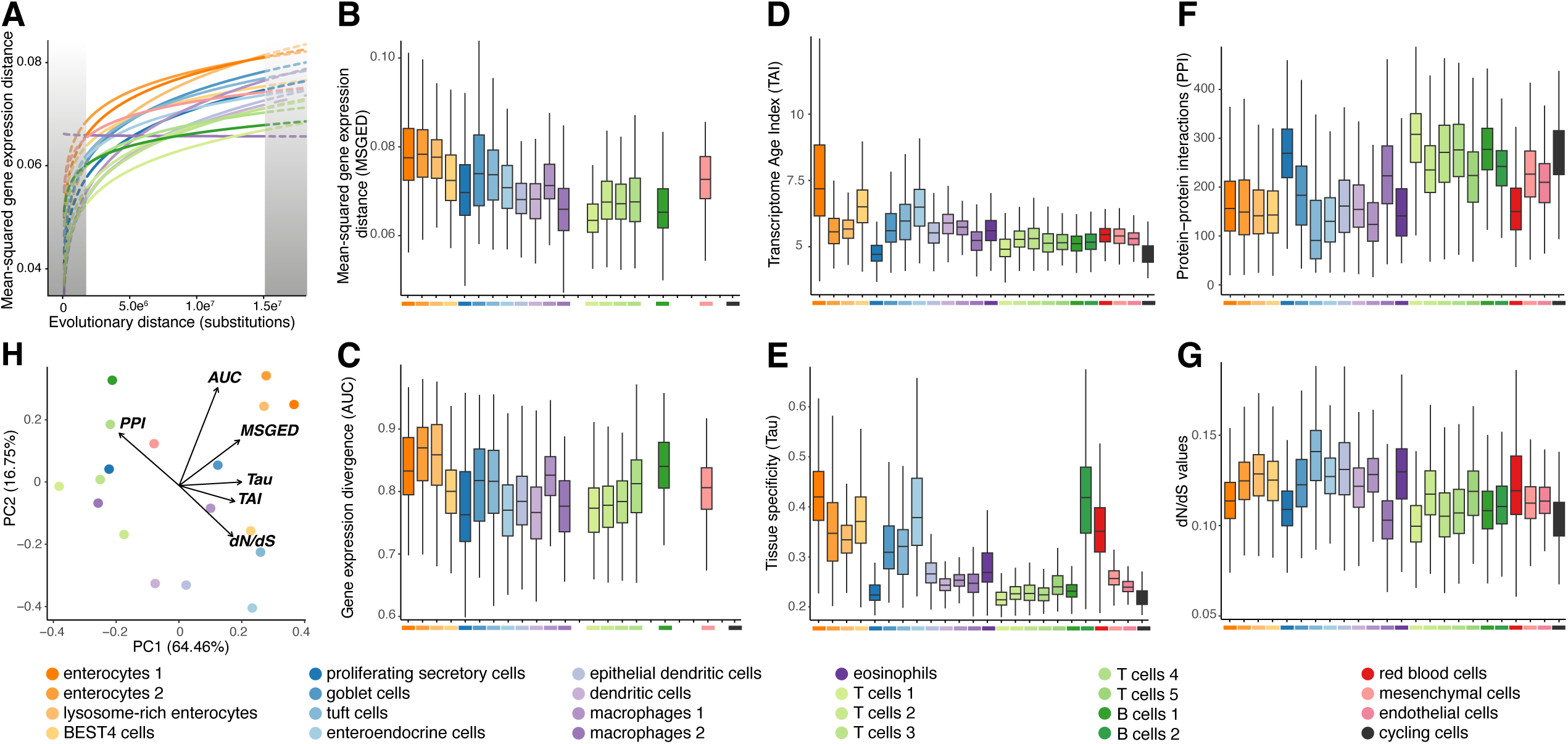
Cell population-specific trajectories of transcriptome evolution in the cichlids’ intestines. (A) Cell population-resolved mean-squared gene expression distances as a function of evolutionary distances (in substitutions), as calculated in EVEE-tools (Chen *et al*., 2019). (B) Mean-squared gene expression distances per cell population. (C) Augur (Skinnider *et al*., 2021) mean cross-validation AUC scores, ranging from 0.5 (random classifier) to 1 (perfect species identification of cells based on gene expression). (D) Transcriptome Age Index (TAI) of expressed genes. (E) Tissue specificity (Tau) scores of expressed genes based on gene expression estimations from 13 cichlid tissues. (F) Predicted number of protein-protein interactions (PPIs) of expressed protein-coding genes, based on the STRING database (Szklarczyk *et al*., 2022). (G) Quantification of signatures of selection in the coding sequences of protein-coding genes, estimated by the ratio of non-synonymous over synonymous substitutions (dN/dS) of expressed genes. Cell populations with a low number of cells or species represented were excluded from analyses shown in panels (A), (B) and (C), resulting in a total of 18 cell populations. (H) Summary principal component analysis of the six evolutionary traits investigated, with the respective loadings indicated by arrows: two independent measures of cell-population-specific cross-species gene expression divergence (MSGED and AUC), and four evolutionary constraints acting on the transcriptome of cell populations (TAI, Tau, PPI, dN/dS), as detailed in panels (B) -(G). See also Figures S1 and S3.

Next, we calculated the phylogenetic age index of each gene (Domazet-Lošo & Tautz, 2010), to then obtain a single transcriptome age index (TAI) for each cell and cell population (Figure 2D). Epithelial cells, and in particular enterocytes 1 – the most anterior absorptive epithelial cells involved in lipid and small molecule metabolism (Figure S2J,K) – displayed higher TAI-values, indicating the expression of ‘evolutionary’ young genes that are possibly under reduced evolutionarily constraints (Murat *et al*., 2023). In contrast, epithelial proliferating cells and cycling cells had the lowest TAI, in line with the conservation of gene sets involved in cellular differentiation (Sun *et al*., 2008; Shay *et al*., 2013) and cell cycle (Cross, Buchler & Skotheim, 2011; Fischer *et al*., 2022). To estimate the extent of pleiotropic constraints in each cell population, we next calculated the tissue-specificity index for each gene, using available bulk RNAseq data from 13 cichlid tissues, namely liver, intestine, lower pharyngeal jaw, gonads, brain, retina, gills, skin, kidney, spleen, heart, blood, and muscle (Brawand *et al*., 2011; Xia *et al*., 2018; Xu *et al*., 2018; Ellison *et al*., 2018; Rajkov *et al*., 2021; El Taher *et al*., 2021; Ricci *et al*., 2023); Figure 2E). Tissue specificity was highest in absorptive epithelial cells and enteroendocrine cells, and low in immune cells, epithelial proliferating cells, mesenchymal cells, and cycling cells, consistent with the broad distribution of the latter cell populations across organs (Rombout *et al*., 2011; Sender *et al*., 2023). As another proxy for pleiotropy, we inferred for each protein-coding gene the number of directly or indirectly inferred high-confidence protein-protein interactions (PPIs) from the STRING database (Szklarczyk *et al*., 2022). We again found divergent patterns in intestinal cell populations, with most epithelial cells, macrophages, and dendritic cells expressing protein-coding genes with overall fewer PPIs than epithelial proliferating cells, T cells, B cells, and mesenchymal cells (Figure 2F). Tissue specificity and putative protein-protein interactions thus suggest that gene expression in epithelial cells, with the exception of proliferating secretory cells, has evolved under reduced pleiotropic constraints in cichlids.

Lastly, to quantify selective pressures acting on the coding sequence of PCGs, we computed dN/dS values for PCGs across distant teleost lineages. We found that the mean dN/dS of epithelial cells is higher than that of the cells of the lamina propria (Wilcoxon test, p < 2.2e^−16^), likely indicating that the coding transcriptome of epithelial cells is evolving under reduced selective pressures. Taken together, our findings suggest that while most intestinal cell populations have evolved under strong evolutionary constraints, intestine-specific absorptive epithelial cells – and in particular enterocytes 1 – show remarkably high cross-species gene expression divergence, in line with relaxed phylogenetic and pleiotropic constraints (Figure 2H, high scores in PC1 and PC2).

### Adaptive divergence in cellular composition of the intestinal epithelium

Changes in abundances of homologous cell types across species have been documented in various tissues (Hunnicutt, Good & Larson, 2022; Johnson *et al*., 2023; Rickelton *et al*., 2024), yet their role in promoting ecological specialization and adaptive phenotypic evolution remains elusive. We therefore explored potential associations between variation in cellular tissue composition in the intestines of Lake Tanganyika cichlids and the different habitats and trophic niches these fishes occupy. Because inter-specific variation in robustness, elasticity, and thickness of the intestines might bias tissue dissociation and, hence, the relative representation of cells of the epithelium versus cells of the underlying lamina propria, we decided to restrict our analyses to epithelial cells only. We found that the relative abundance of epithelial cell populations varied substantially across species, in particular with respect to the relative proportions of absorptive to non-absorptive cells (Figure 3A). Using linear regression (lm) and phylogenetic generalized least squares (pGLS) analyses, we then tested for associations between relative epithelial cell population abundances and a set of previously established proxies for the foraging ecology of Lake Tanganyika cichlids (stable carbon isotope signatures [δ^13^C-value] as a proxy for the benthic-pelagic habitat axis, PC1 of oral jaw morphology) as well as for their dietary niches (stable nitrogen isotope signatures [δ^15^N-values] as proxy for the trophic level (Figure 3A), normalized gut length, PC2 of lower pharyngeal jaw [LPJ] morphology and a diet ecology vector that summarizes these three dietary predictors) ((Ronco *et al*., 2021; Duque-Correa *et al*., 2024); Figure 3B-D, Figure S4A-C and Methods). We found strong positive correlations between the relative abundance of absorptive epithelial cells and δ^15^N-values across species (Figure 3D, lm and pGLS: p = 7e^−4^, R^2^ = 0.43, λ = 0). This trend is largely driven by an overrepresentation of enterocytes 1 in the guts of cichlid species at higher trophic levels, e.g. piscivores and scale eaters, and a relative underrepresentation of enterocytes 1 in species feeding on algae and plants (Figure 3B,D). We observed a similar pattern in our spatial deconvolution approach, which revealed higher abundance of enterocytes 1 in the two most anterior gut sections of the piscivore *Lepidiolamprologus attenuatus* (Lepatt) compared to the two other species *Eretmodus cyanostictus* (Erecya) and *Astatotilapia burtoni* (Astbur) that feed on plants, diatoms, and small arthropods (Figure S2C). By contrast, proliferating secretory cells (Figure 3C,D, lm and pGLS: p = 3e^−3^, R^2^ = 0.35, λ = 0), goblet cells, enteroendocrine cells, and epithelial dendritic cells all showed negative correlations of abundances with δ^15^N-values, indicating that these cell types are more numerous in the guts of primary consumers (Figure 3D, lm and pGLS: R^2^ > 0.2, p < 0.05) than in species at higher trophic levels (Figure 3D, lm and pGLS: p = 3e^−3^, R^2^ = 0.35, λ = 0). Taken together, our results indicate that the relative cellular abundance of the most anterior intestinal epithelial absorptive cells may indeed represent an adaptive response to novel dietary niches, potentially facilitating changes in lipid metabolism and small molecule transport (Figure S2F,G), but not in more posterior enterocytes. More importantly, this suggests that also at the cellular level of the intestinal tissue, there is a strong phenotype-environment correlation in the cichlid adaptive radiation in Lake Tanganyika.

**Figure 3.**
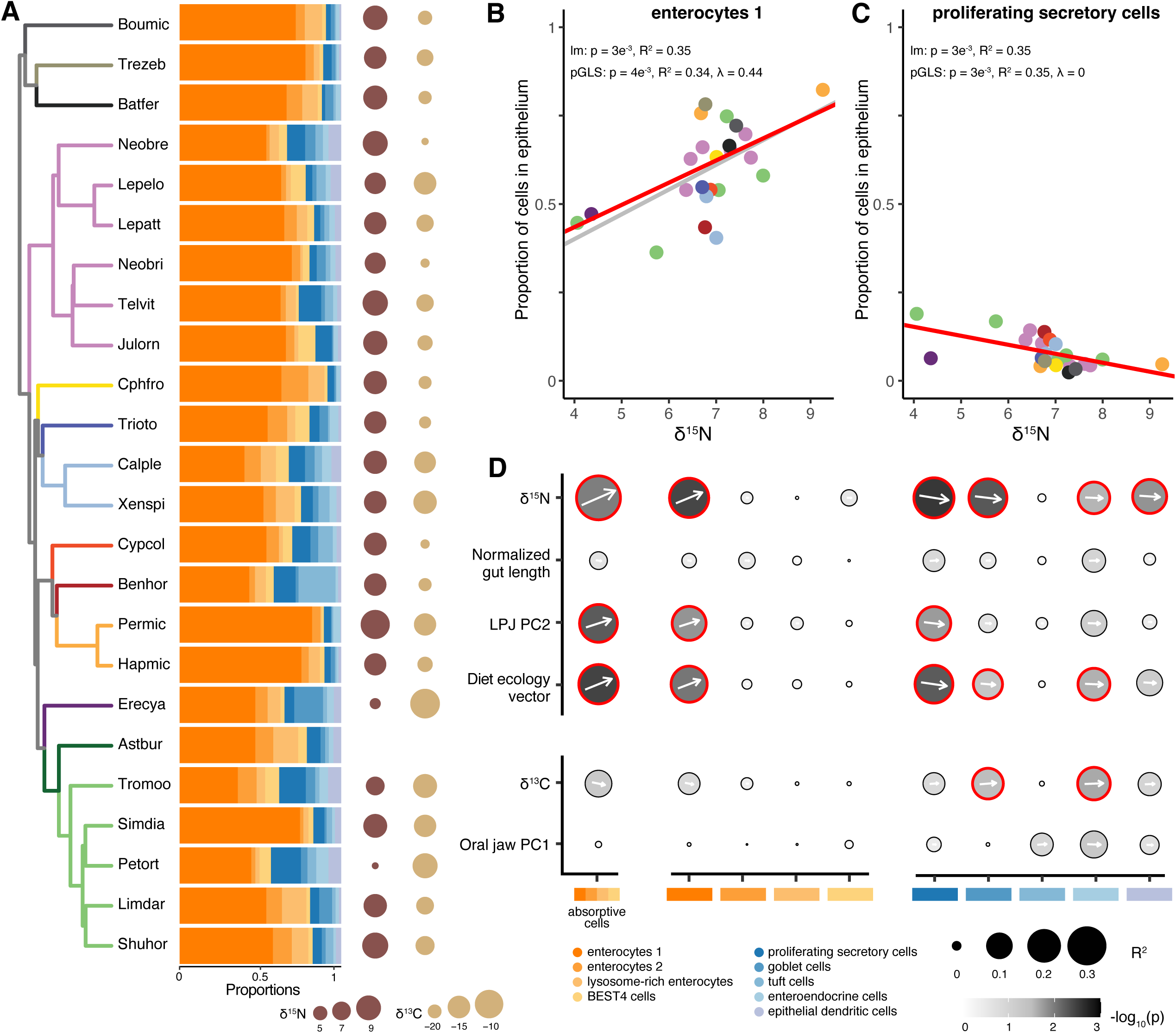
Epithelial cell population proportions are associated with dietary specialization. (A) Left: Time-calibrated genome-wide species tree of the 24 cichlid species from Lake Tanganyika included in this study. Branch colors refer to the cichlid tribes as detailed in Ronco and colleagues (Ronco *et al*., 2021). Middle: Relative abundance of absorptive (orange) and non-absorptive (blue) cell populations in the intestinal epithelium of each cichlid species. Right: Nitrogen stable isotope (δ^15^N-value) and carbon stable isotope (δ^13^C-value) signatures estimated for each species (Ronco *et al*., 2021). (B) and (C) Linear model (lm) and phylogenetic generalized least squares (pGLS) analyses showing a strong correlation between δ^15^N-values and the relative abundance of enterocytes 1 (B), and proliferating secretory cells (C), respectively. The colors of the points in panels (B) and (C) again refer to the different cichlid tribes as in (A). (D) Summary dot plot of phylogenetic regressions (pGLS) results between six ecological predictors, indicative of diet (top) or foraging ecology (bottom), and the relative abundance of epithelial cell populations. Statistically significant correlations (p < 0.05) are indicated with a red outline, and the arrows indicate the direction of the correlation (arrows pointing upwards and downwards indicate positive and negative correlations, respectively). See also Figure S4.

### Ecological signals in gene expression profiles of intestinal cell populations

To further characterize the molecular and cellular responses to ecological niche adaptations beyond cellular variations in intestinal epithelium composition, we examined the association between gene expression and ecological predictors. For each cell population, we performed multivariate phylogenetic regressions (Clavel, Escarguel & Merceron, 2015); see Methods for details) in three sets of genes, representing broadly expressed genes in at least 20% of cells (Figure 4A), cell-population-specific genes (Figure 4B), and genes co-expressed in scWGCNA modules (Figure 4C), respectively. In the ‘global’ dataset of broadly expressed genes, ecological associations largely reflected overall gene expression affinities across cell populations, with most of the epithelial cell populations showing similar correlations with ecological predictors (Figure 4A). Analyses of more restricted gene sets revealed stronger ecological signals within some specific epithelial cells (Figure 4B,C). In particular, enteroendocrine cells displayed strong signals of associations with both foraging ecology and diet (Figure 4A-C), while enterocytes 1 displayed strong significant associations of gene expression signatures with several dietary predictors (δ^15^N-values, normalized gut length, and diet ecology vector), in all three sets of genes (Figure 4A-C). This dietary signature was confirmed by a two-block partial least-squares (2B-PLS) analysis, which revealed that δ^15^N-values and normalized gut length were the primary ecological predictors of gene expression variation in enterocytes 1 (Figure S4D-F; Wilcoxon text, p < 4e^−4^). In contrast, other enterocyte populations showed weaker and mostly ‘global’ dataset-specific associations with ecological factors, indicating a less pronounced cell population-specific involvement in adaptations to different dietary niches. These findings again emphasize the pivotal role of enterocytes 1 in dietary adaptations in cichlids. Furthermore, four scWGCNA modules primarily expressed in enterocytes 1 showed strong diet-related signals (modules 7, 9, 16, and 17 in Figure 4C). Notably, these modules were found to be enriched in genes coding for transporter, transmembrane and solute carrier proteins, as well as in factors involved in amino acid absorption and peptide digestion (Figure S1J,K), highlighting the importance of these functions in cellular adaptations to novel dietary niches, and assigning them to a particular cell population.

**Figure 4.**
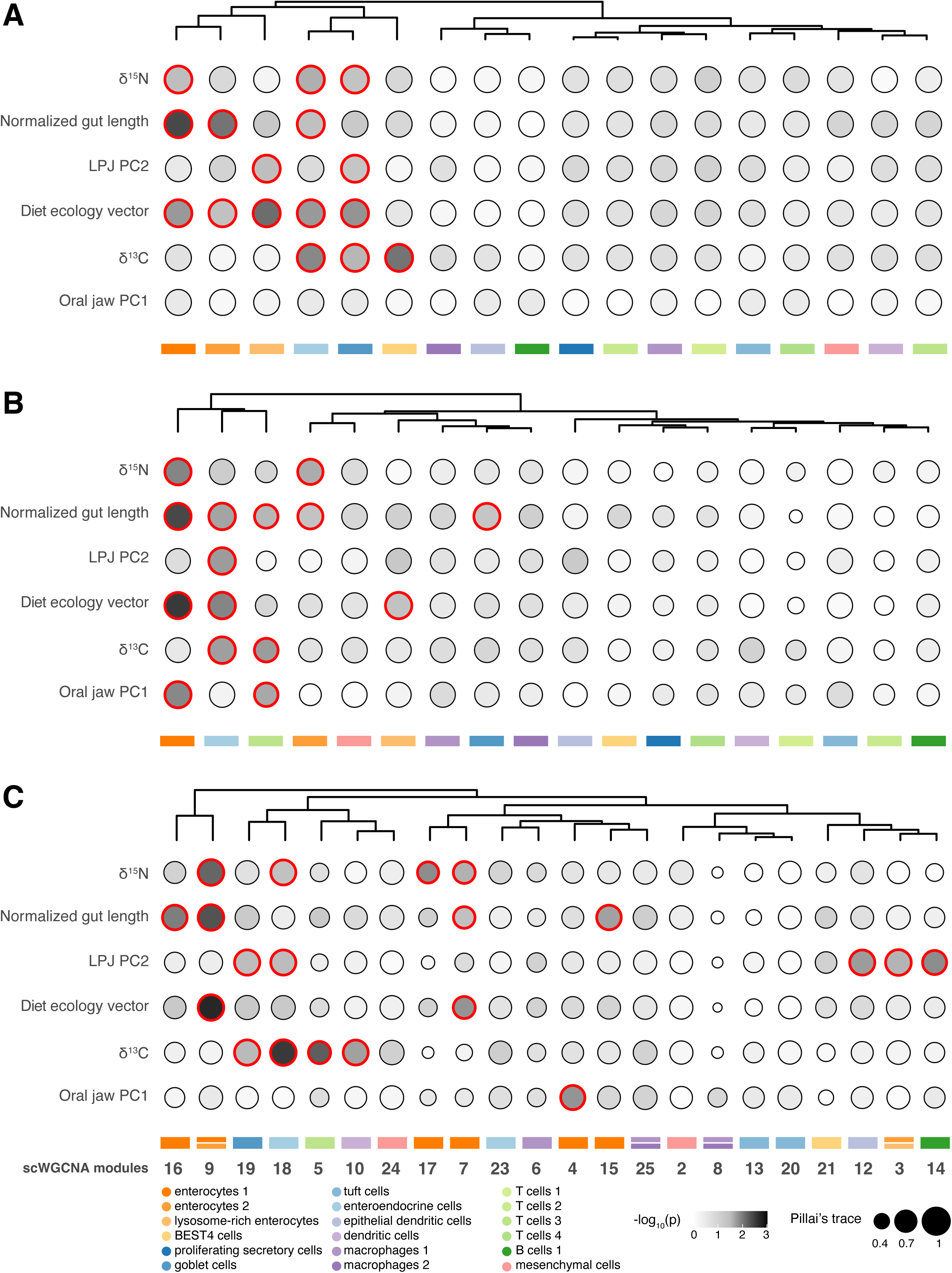
Correlations between gene expression signatures in intestinal cell populations and ecology. (A), (B) and (C) Summary dot plots and hierarchical clustering of multivariate phylogenetic regressions (type II phylogenetic MANOVA, computed with mvMORPH) between six ecological predictors and gene expression in cell populations based on (A) genes expressed in at least 20% of the cells within each cell population, (B) cell population-specific genes, and (C) genes in scWGCNA co-expression modules. Size of circles indicate Pillai’s trace values (Engel *et al*., 2015), with statistically significant correlations (p<0.05) indicated with a red outline, and hierarchical clustering based on -log10(p). See also Figure S4.

### Molecular basis of dietary adaptations in anterior enterocytes

To uncover the molecular underpinnings of intestinal adaptations at the cellular level in cichlid fishes from Lake Tanganyika, we identified genes showing strong expression associations with ecological predictors in enterocytes 1, the cell population with the strongest ecological signals in both relative abundance and gene expression, using the intersection of four phylogenetic comparative approaches

(Methods, Figure S5A,B). Consistent with the strong dietary signature in relative abundance and gene expression of this cell population, we found between 23 and 98 genes associated with each dietary predictor (δ^15^N-values, normalized gut length and diet ecology vector). Interestingly, we also detected 17 genes correlating with δ^13^C-values, likely due to the interdependence of nitrogen and carbon stable isotope signatures, as certain food sources are restricted to specific macrohabitats. We then assessed the tissue- and cell-population-specificities of these genes by comparing them to all expressed genes in enterocytes 1, regardless of their ecological relevance. Genes correlating with one or several ecological predictors exhibited significantly higher tissue specificity (Figure 5A; Wilcoxon test, p < 0.001) and gene specificity indices (Figure 5B; Wilcoxon test, p < 0.05 for 5/6 gene sets) than the average of all expressed genes (referred to as ‘Universe’ in Figure 5A-C). These findings indicate reduced pleiotropy and a higher potential for adaptive expression changes, as previously speculated in other freshwater fish (Papakostas *et al*., 2014). Further, we explored whether these gene specificities were associated with higher rates of gene expression evolution. By modeling gene expression through a Brownian motion (BM) process, we evaluated the rate of evolution of each expressed gene using the σ^2^-parameter (Harmon *et al*., 2008). Our analyses revealed that ecologically relevant genes evolved at significantly higher rates than the remaining genes (Figure 5C; Wilcoxon test, p < 0.001). This suggests that regulatory evolution in these genes has been both adaptive and exceptionally fast, driving the functional and molecular specializations that enable rapid diversification into distinct ecological niches. To further investigate the rapidly evolving, enterocytes 1-specific genes, we performed GO enrichment analysis, focusing on genes with expression patterns correlating either positively or negatively with trophic level (δ^15^N-values; Figure S5C,D). Genes positively correlated with trophic level were primarily involved in ion and lipid binding, peptidase activity, catabolic processes, and cellular structure, hinting at the importance of efficient breakdown and absorption of fats and proteins in species at higher trophic levels. In contrast, genes negatively correlated with trophic level were linked to cell adhesion, tyrosine kinase activity, lipid metabolism, ion transport, tube development, and signaling pathways. This may reflect adaptations for a rapid turnover and maintenance of epithelial cells, to preserve gut integrity, and to facilitate efficient ion balance, as is typical for herbivorous organisms feeding on mineral-rich plant-based diets (Bakke-McKellep *et al*., 2007; He *et al*., 2015; Pelster *et al*., 2015). Overall, these patterns suggest a strong connection between gene function, expression levels, and adaptations to nutrient absorption and uptake.

**Figure 5.**
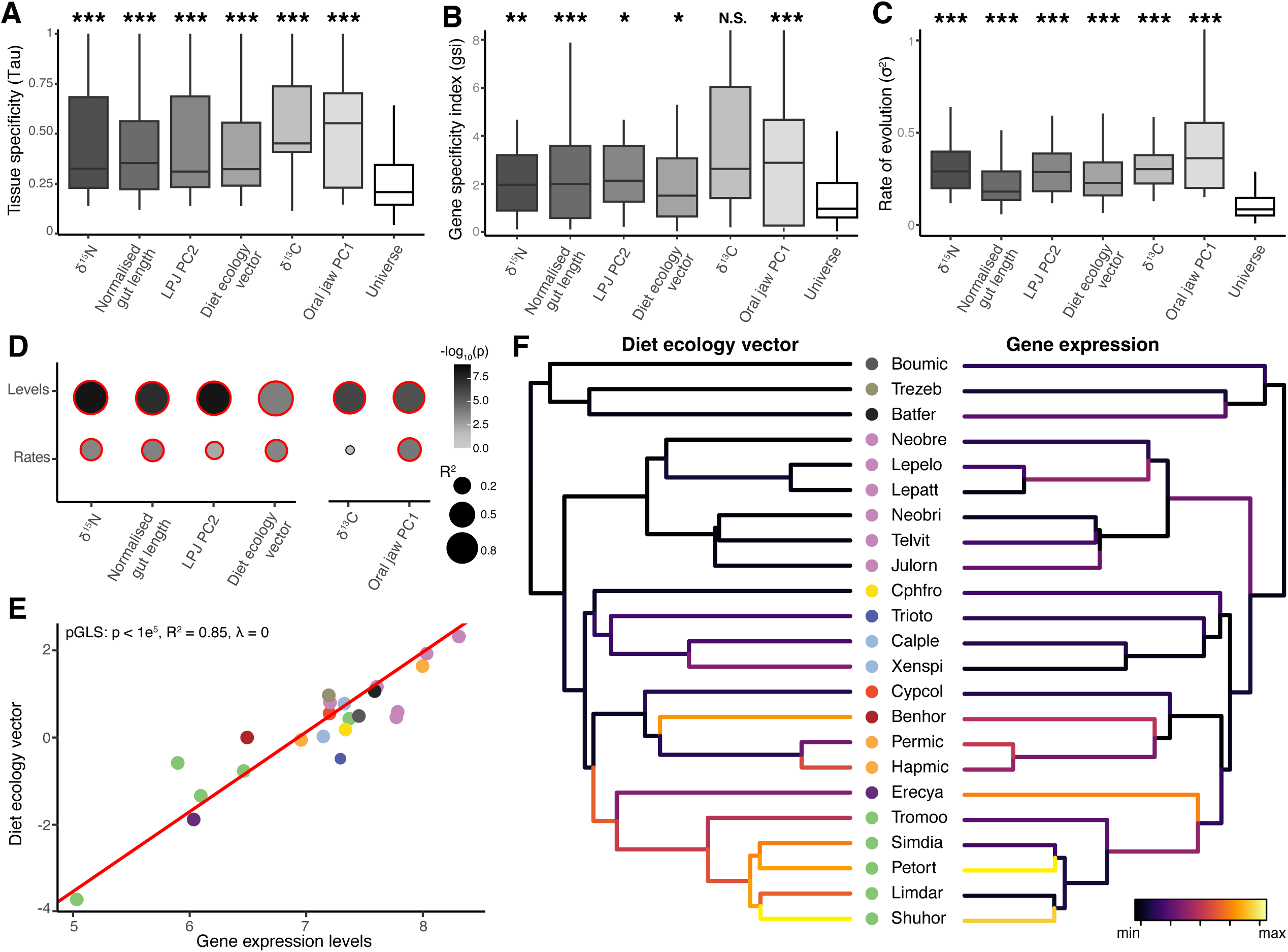
Gene expression evolution of diet-related genes in anterior enterocytes. (A) Tissue specificity scores (Tau) of genes whose expression is associated with ecological predictors in enterocytes 1. (B) Gene specificity indices (gsi) of genes whose expression is associated with ecological predictors in enterocytes 1. (C) Rate of gene expression evolution (σ^2^) inferred from a Brownian motion process for genes whose expression is associated with ecological predictors in enterocytes 1. In panels (A), (B) and (C), statistical difference between the ‘Universe’ genes and each gene set was assessed by a Wilcoxon test. ***: p < 0.001, **: p < 0.01, *: p < 0.05, N.S.: non-significant. (D) Top: summary dot plot of correlations between six ecological predictor values and species-specific gene expression levels of ecologically relevant genes. Bottom: summary dot plot of correlations between the branch-specific evolutionary rates of the six ecological predictors and the branch-specific evolutionary rates of gene expression of ecologically relevant genes, across the phylogeny. Statistically significant correlations (p < 0.05) are indicated with a red outline. (E) Phylogenetic generalized least squares (pGLS) analyses showing strong, significant correlation between the diet ecology vector and mean species-specific expression levels of genes associated with the diet ecology vector. The colors represent the cichlid tribes of the species, as in Figure 3A. (F) Time-calibrated species tree with branches colored according to the branch-specific evolutionary rates for the diet ecology vector (left), and for the expression of genes associated with the diet ecology vector (right). See also Figures S4 and S5.

Finally, we explored how these genes may have contributed to the rapid ecological diversification of cichlid fishes in Lake Tanganyika. We first assessed the strength of association between various ecological predictors and species-specific mean gene expression levels across each gene set. We identified strong associations for both diet and foraging ecology predictors (Figure 5D, top). For example, we found that enterocyte 1 gene expression profiles correlate strongly with the diet ecology vector (Figure 5D,E; lm and pGLS: R² = 0.85, p < 1e^5^, λ = 0). We then inferred evolutionary rates for both ecological predictors and gene expression along the species phylogeny, using data from 126 – 234 cichlid species in Lake Tanganyika for ecological predictors and scRNAseq data from 23 species for gene expression (Figure 5D, bottom). Despite the stark difference in species numbers and species sampling density along the cichlid phylogeny, the evolutionary rates of dietary predictors – particularly dietary ones – and of the associated relevant gene sets showed highly significant correlations (lm: p < 0.005, R^2^ > 0.17), except for our habitat proxy (δ^13^C-values). Interestingly, we detected heterogeneous rates of evolution across branches and identified accelerated evolution in both gene expression and predictors in the Tropheini (Tromoo, Simdia, Petort, Limdar, Shuhor in Figure 5F, Figure S5E-G), a tribe known for its dietary transitions to algae browsing, algae grazing and even piscivory (Singh *et al*., 2022). Our trait evolutionary analysis also confirmed previously identified pulses of ecological diversification (Ronco *et al*., 2021), including a major pulse of dietary diversification around 1.5-4 million years ago (Figure S5H, asterisks; δ^15^N-values, normalized gut length, PC2 of LPJ morphology, diet ecology vector), which likely reflects fine-scale ecological niche partitioning through trophic specialization (Ronco *et al*., 2021; Ronco & Salzburger, 2021).

Collectively, our findings therefore suggest that adaptations in gene expression regulation within anterior enterocytes have contributed to feeding specializations, leading to rapid trophic niche partitioning and accelerated species diversification in the later stages of the adaptive radiation of cichlid fishes in Lake Tanganyika.

## Discussion

In this study, we generated extensive single-cell RNA sequencing data from the intestines of 24 ecologically and morphologically highly diverse species of cichlid fishes from Lake Tanganyika to investigate the cellular and molecular basis of dietary adaptations in the digestive tracts of these rapidly evolving species. We first show that, overall, the epithelial and immune cell type diversity in the intestinal tissue of cichlid fishes is comparable to that of other vertebrate intestines (Elmentaite *et al*., 2021), despite extensive morphological and physiological diversification (Stevens, 1990); Figure 1). We further found that the different absorptive epithelial cell populations are not evenly distributed along the anterior-posterior axis of the cichlids’ intestines, with lysosome-rich enterocytes (LREs) – previously only identified in stomachless fishes (Park *et al*., 2019) – being more abundant in the posterior part of the intestine, and a transcriptionally distinct population of enterocytes, here referred to as enterocytes 1, in the anterior part (Figure S2).

By combining our multi-species gene expression data at cellular resolution with comprehensive ecological and morphological information, we demonstrate that, out of all the epithelial cell types, enterocytes 1 exhibit the strongest signals of dietary adaptations with respect to both relative cellular abundances and gene expression profiles. Interestingly, these enterocytes are transcriptionally most similar to mouse duodenal enterocytes (Lickwar *et al*., 2017), which have been shown to mediate lipid metabolism (Zwick *et al*., 2024) and to display rapid plastic responses to novel food sources (Clara *et al*., 2017; Enriquez *et al*., 2022). In cichlids, enterocytes 1 are overrepresented in carnivorous species compared to primary consumers, which in turn feature the highest relative proportions of secretory cells (Figure 3B-D). Our results are thus in line with previous plasticity experiments revealing increased numbers of absorptive cells in mice fed a high-fat diet (Enriquez *et al*., 2022), and of goblet cells and proliferating cells in fish and mammals fed on high-starch (Huang *et al*., 2021), or fiber-rich diets (Ito *et al*., 2009; Hino *et al*., 2012; Saqui-Salces *et al*., 2017), but, here in an evolutionary-comparative context. The diet-related divergence in tissue composition detected among the different cichlid species likely reflects the distinct physiological and tissue-level requirements for dietary specialization. Herbivorous and algivorous species that consume diets rich in fibers require more mucus-secreting cells – which can contribute to plant nutrient extraction (Tibbetts, 1997) – and a more rapid renewal of secretory cells in general. In contrast, carnivorous species that feed on fat-rich diets may rely on a more efficient lipid metabolism and nutrient absorption along their shorter intestines. Since both absorptive and secretory cell lineages in the intestinal epithelium originate from a common pool of stem cells, our results indicate that epithelial cell fate specification programs (Crosnier *et al*., 2005; Fre *et al*., 2011) or progenitor pool expansion (Tóth *et al*., 2017) can be ecologically tuned. This further suggests that, at least in cichlids, shifting the ratio between absorptive versus secretory cells is a major axis of rapid adaptation and specialization to diverse nutritional sources.

At the molecular level, we found strong associations between dietary niche and gene expression signatures in anterior enterocytes (Figure 4). We further show that these associations are primarily driven by tissue- and cell-population-specific, and fast-evolving genes (Figure 5A-C) likely evolving under positive selection and lower evolutionary constraints (Wittkopp & Kalay, 2011; Wang, Starr & Fraser, 2024). This partial relaxation of evolutionary and pleiotropic constraints, which has been previously linked to overall lower functional constraints and to adaptive evolution (Hodgkin, 1998; Carroll, 2005; Cardoso-Moreira *et al*., 2019; Zhang, 2023), may have enhanced the potential of cell-population-specific genes to be more dynamically regulated and fine-tune their expression to changing physiological demands in response to novel food sources. This is further evidenced by the high overall interspecific gene expression divergence observed in anterior enterocytes (Figure 2A-C), and by the increased rates of gene expression evolution of diet-associated genes in specific branches of the species tree, which mirror the documented rapid trophic specializations and transitions in the later phase of the cichlid radiation in Lake Tanganyika (Wagner *et al*., 2009; Muschick *et al*., 2012; Ronco *et al*., 2021; Singh *et al*., 2022); Figure 5D-F; Figure S5E-H).

In summary, we provide evidence that trophic specializations in the digestive tract of cichlid fishes have occurred across multiple layers of biological organization, ranging from macro-morphological changes to cellular and molecular adaptations at the tissue-and transcriptome-level of their intestines. Furthermore, our study demonstrates the power of integrating single-cell transcriptomics with morphological and ecological data, to help uncover the complex cellular and molecular underpinnings of phenotypic evolution and organismal diversification within the context of adaptive radiations.

## Data and code availability

All single-cell and bulk RNA-seq raw and processed data generated in this study have been deposited at GEO (https://www.ncbi.nlm.hiv.gov/geo/) under accession numbers GSE280410 and GSE280411. All original code is available on GitHub (https://github.com/AntoineFages/scRNA_paper).

Any additional information required to reanalyze the data reported in this paper is available from the lead contact upon request.

## Acknowledgements

We thank Heinz Büscher, Julia Barth, and Frédéric Schedel for providing fish specimens, Attila Rüegg for fish care, Julia Johnson for fish illustrations, Nicolas Feigenwinter, Sabrina Fischer, and Nicolas Boileau for assistance in laboratory work. We further thank Marcus Clauss and María José Duque-Correa for discussions on fish diets and comparative physiology; Fabio Sacher, Xuefei Yuan, and Leticia Rodríguez Montes for single-cell bioinformatic support; Prisca Liberali for discussions on intestinal stem cell and progenitor biology, Henrik Kaessmann for valuable discussions on cell type and tissue evolution; and the Genomics Facility Basel of the ETH Zurich Department of Biosystems Science and Engineering for next-generation sequencing. Calculations were performed at sciCORE (http://scicore.unibas.ch/) scientific computing core facility at University of Basel. This work was supported by the Swiss National Science Foundation (SNSF) Sinergia grant n. 189970.

## Author Contributions

Conceptualization: A.F., W.S., P.T.; formal analysis: A.F., F.R.; investigation: A.F., M.L., C.E.T.H., P.N.K.M.; resources: A.I., W.S., P.T.; writing - original draft: A.F., W.S., P.T.; writing - review and editing: A.F., M.L., F.R., C.E.T.H., N.M., A.I., W.S., P.T.; visualization: A.F., F.R., W.S., P.T.; funding acquisition: W.S., P.T.

## Supplemental figure

**Figure S1.**
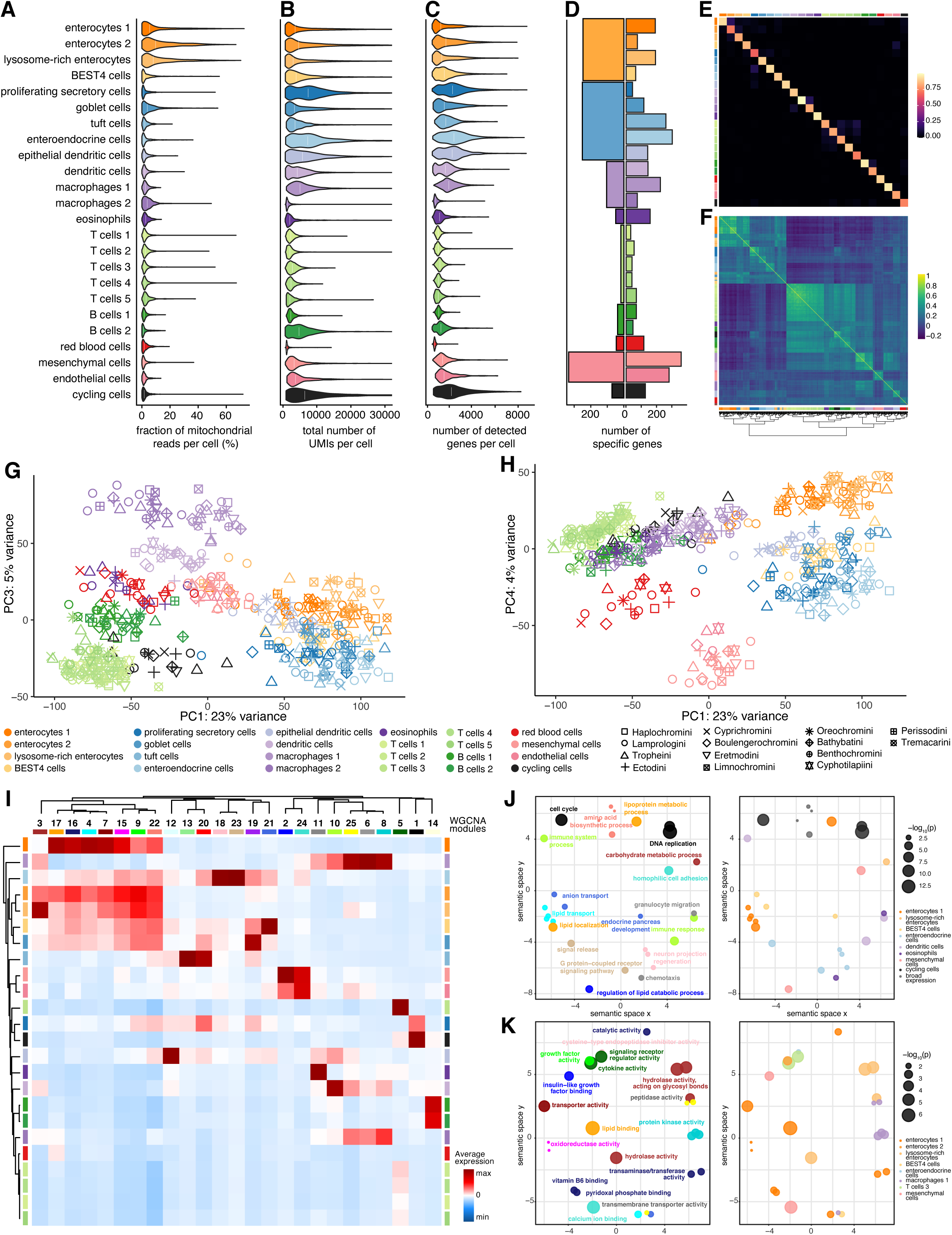
Characterization of intestinal cell populations, related to Figure 1 and Figure 2. (A) Fraction of reads mapping to mitochondrial genes in intestinal cell populations (means=2-14%). (B) Total number of unique molecular identifiers (UMIs) detected per cell in intestinal cell populations (means=861-9157). (C) Number of detected genes per cell in intestinal cell populations (means=347-2148). (D) Number of specific genes detected in intestinal cell populations (top) and cell type categories (down). (E) Confusion heatmap of cross-validation results for the 24 distinct cell populations identified in the cichlid intestine. The gradient scale indicates the proportion of correct cross-validations, and ranges from 0 to 1. Row and column annotation colors represent the intestinal cell populations. (F) Pairwise Spearman’s rank correlation matrix with hierarchical clustering of pseudobulks for each cell population and species in the dataset, ranging from 0 to 1. Row and column annotation colors represent the intestinal cell populations. (G) PC1 and PC3 of PCA of pseudobulk gene expression variation across all cell populations and species. Colors refer to cell populations and shapes to the tribes to which the species belong. (H) PC1 and PC4 of PCA of pseudobulk gene expression variation across all cell populations. (I) Heatmap of average scWGCNA module eigengene expression across intestinal cell populations, scaled across scWGCNA modules, with associated hierarchical clusterings of cell populations and scWGCNA modules. Column annotation colors represent the scWGCNA modules, row annotation colors represent the intestinal cell populations. (J) REVIGO scatterplot visualization of Biological Process gene ontology (GO) terms enriched in scWGCNA modules, colored by scWGCNA modules (left) or cell population of highest expression for each module (right). (K) REVIGO scatterplot visualization of Molecular Function gene ontology (GO) terms enriched in scWGCNA modules, colored by scWGCNA modules (left) or cell population of highest expression for each module (right). See also Table S2.

**Figure S2.**
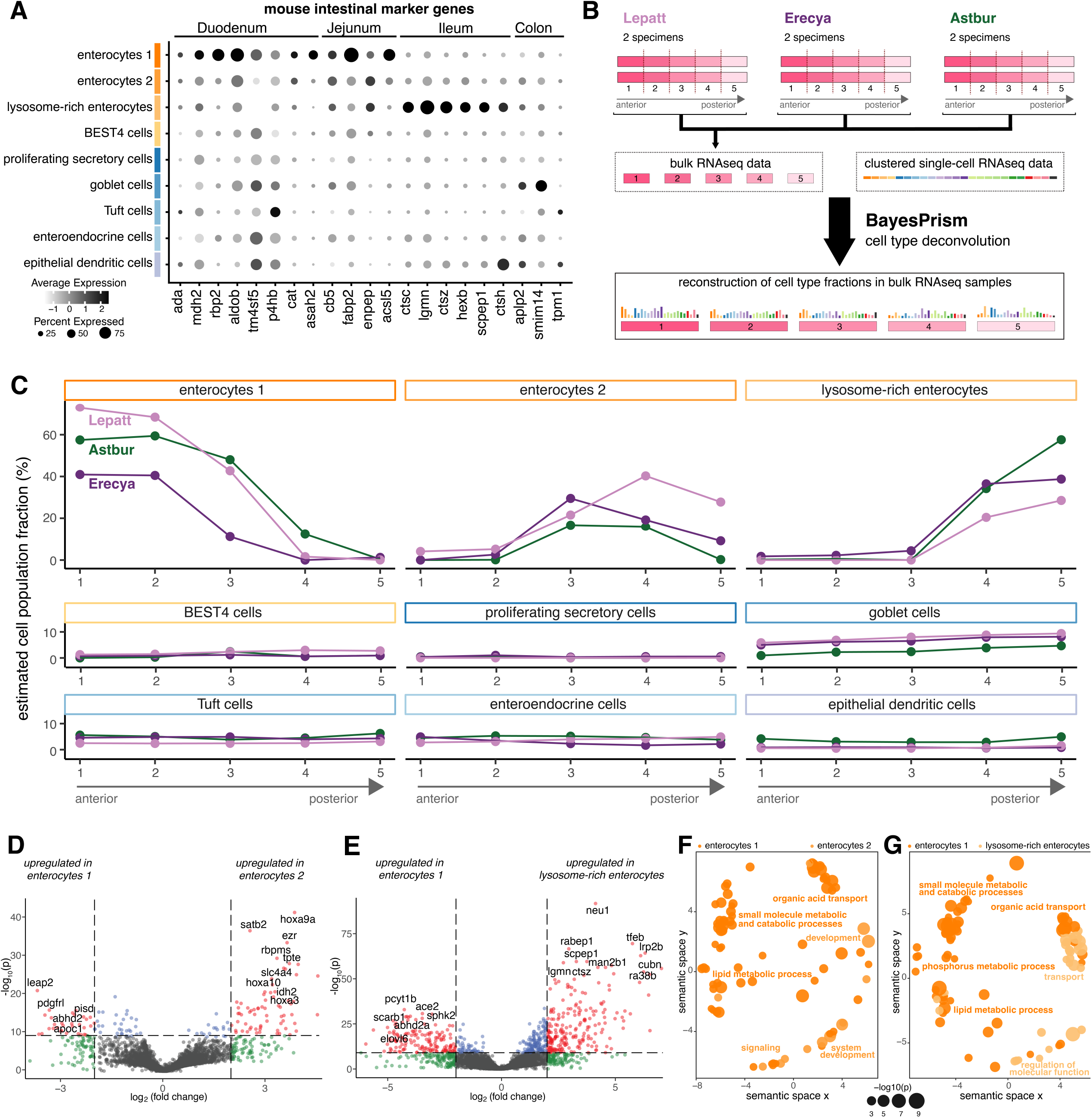
Anterior-posterior characterization of epithelial cell populations, related to Figure 1. (A) Dot plot of expression of mouse intestinal marker genes specific to duodenum, jejunum, ileum and colon, as identified by Lickwar and colleagues (Lickwar *et al*., 2017). (B) Experimental design and workflow of cell type deconvolution of cichlid intestine bulk RNAseq data informed by scRNAseq data, using BayesPrism (Chu *et al*., 2022). (C) Estimated cell population fractions in the five intestinal parts of three African cichlid species. The colors indicate the species’ tribe colors used in Figure 3. (D) , (E) Volcano plots of differential expression analysis between enterocytes 1 and enterocytes 2 (D), and enterocytes 1 and lysosome-rich enterocytes (E). Red dots indicate differentially expressed genes. (F), (G) REVIGO scatterplot visualizations of Biological Process GO terms enriched in differentially expressed genes between enterocytes 1 and enterocytes 2 (F), or in differentially expressed genes between enterocytes 1 and lysosome-rich enterocytes (G). See also Table S3.

**Figure S3.**
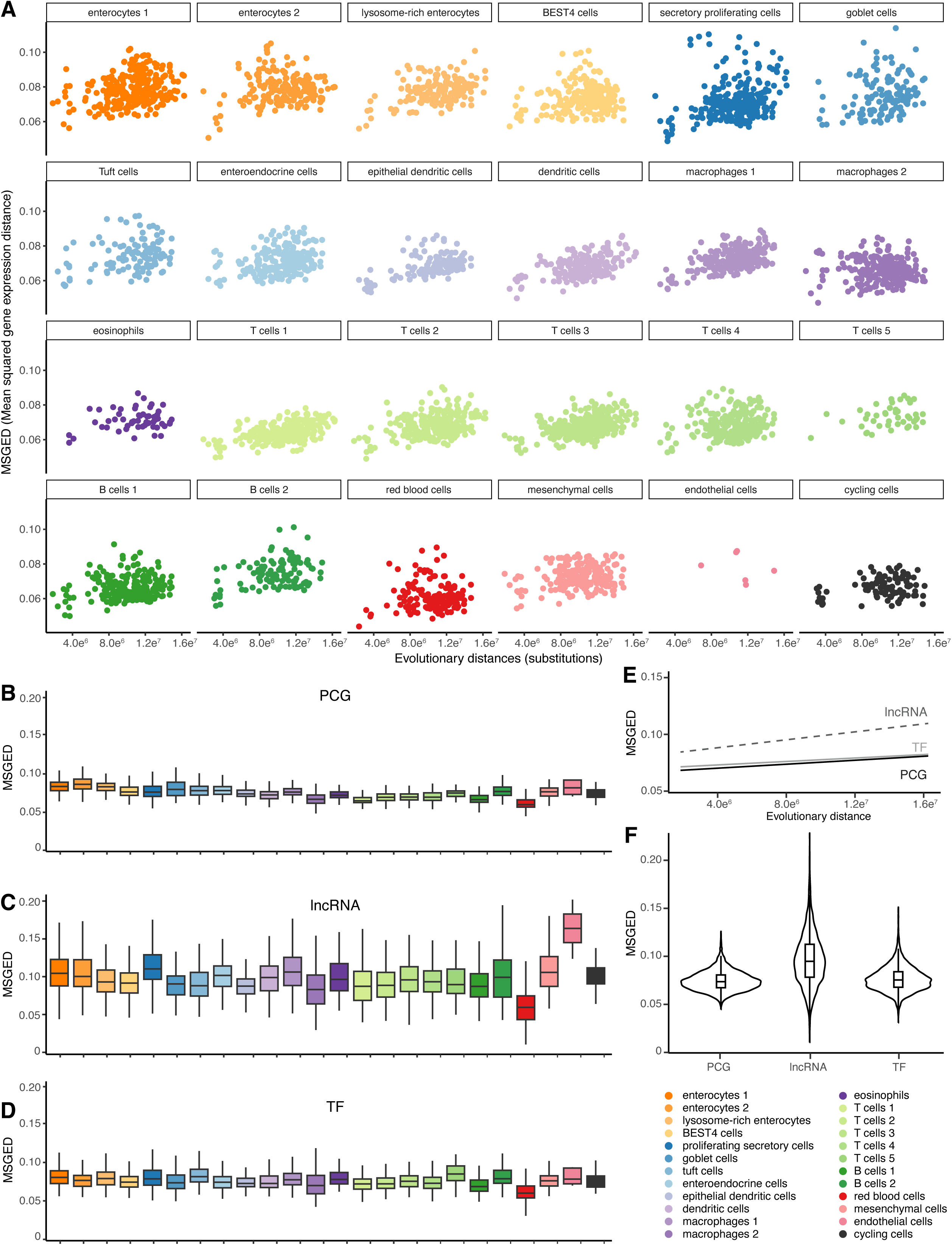
Cross-species gene expression divergence in intestinal cell populations, related to Figure 2. (A) Mean-squared gene expression distances along evolutionary distances (measured in genomic substitutions) calculated across pairs of species in intestinal cell populations. (B) Average mean-squared gene expression distances calculated across pairs of species and based only on protein-coding genes (PCGs) only, in each intestinal cell population. (C) Average mean-squared gene expression distances calculated across pairs of species and based only on long non-coding RNAs (lncRNAs) only, in each intestinal cell population. (D) Average mean-squared gene expression distances calculated across pairs of species and based only on transcription factors (TFs) only, in each intestinal cell population. (E) Mean-squared gene expression distances along evolutionary distances (measured in genomic substitutions) calculated across pairs of species, based on expression of PCGs (dark gray solid line), lncRNAs (gray dashed line) and TFs (light gray solid line). (F) Average mean-squared gene expression distances calculated across pairs of species and based on expression of PCGs, lncRNAs and TFs.

**Figure S4.**
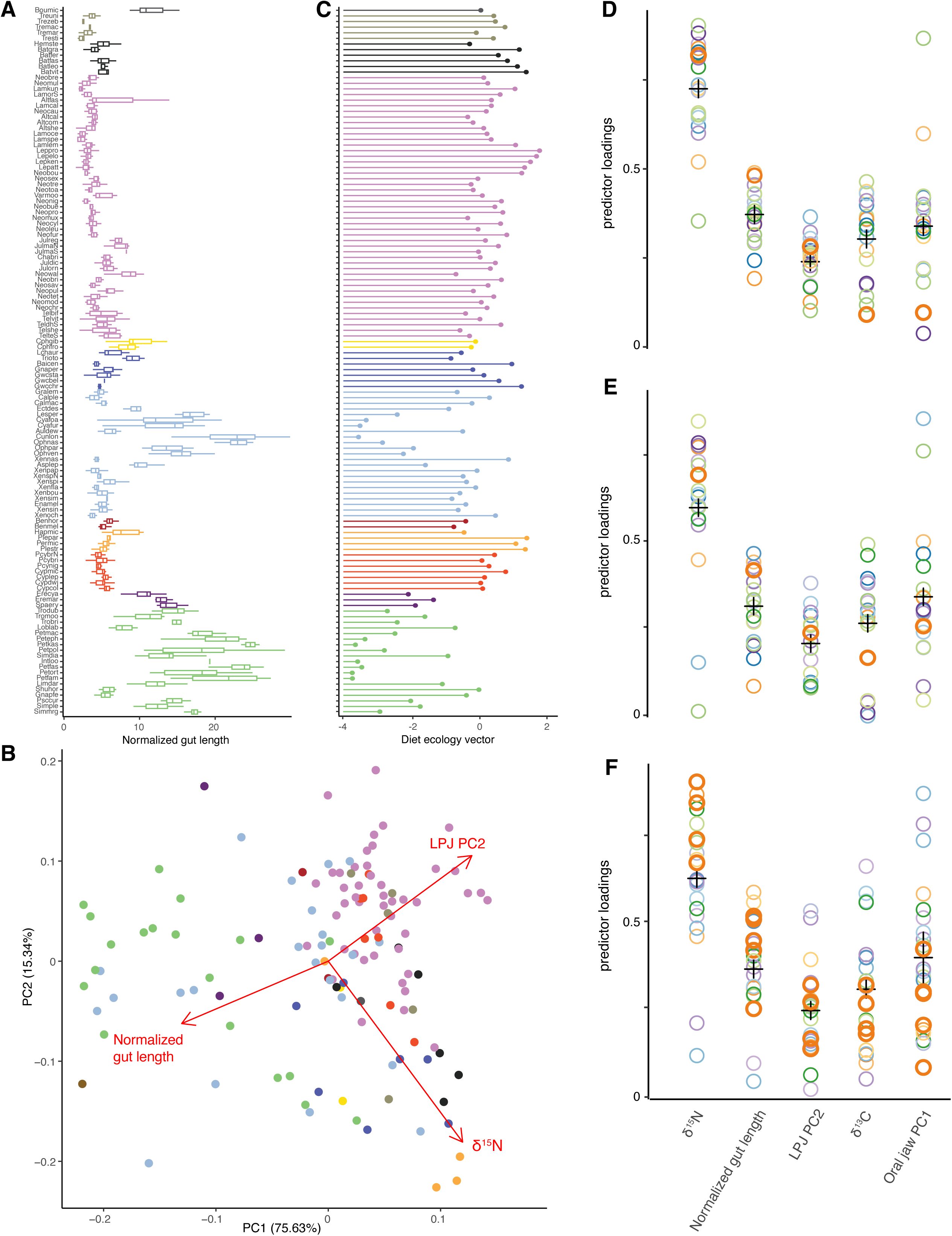
Ecological predictors and their associations with intestinal epithelium gene expression signatures, related to Figures 3-5. (A) Normalized gut length, calculated as the ratio of total gut length to the cubic root of the weight measured in wild-caught specimens of 126 Lake Tanganyika cichlid species. (B) PCA based on three dietary predictors (δ^15^N, normalized gut length and PC2 of LPJ morphology) measured across 126 Lake Tanganyika cichlid species. (C) PC1 of the PCA shown in panel (B), used as diet ecology predictor in ecological association analyses. (D) , (E) and (F) Ecological predictor loadings of a phylogenetic two-block partial least-squares (p2B-PLS) analysis of ecological predictors and cell population gene expression values. (D) is based on genes expressed in at least 20% of the cells within each cell population, (E) on cell population-specific genes, and (F) on genes in scWGCNA co-expression modules.

**Figure S5.**
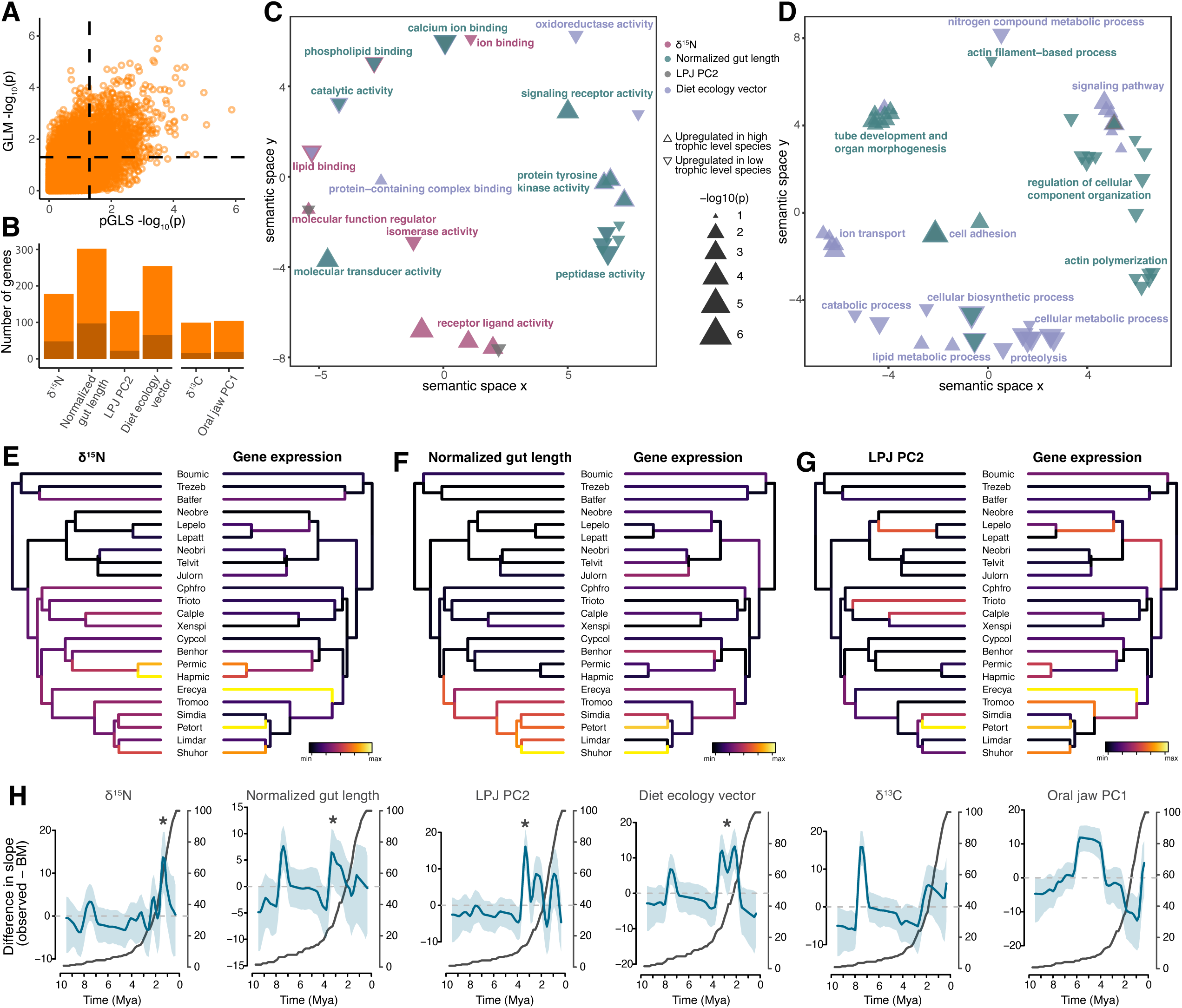
Characterization and evolution of ecologically-relevant genes in anterior enterocytes, related to Figure 5. (A) Distributions of p-values from lm and pGLS analyses, measuring the strength of association between gene expression and ecological predictors in enterocytes 1. (B) Intersection of the four approaches used for finding genes whose gene expression correlates with ecological predictors. Lighter orange represents the number of genes identified through univariate approaches, and darker orange the strong candidate genes, identified as intersections of univariate and multivariate approaches. (C) REVIGO scatterplot representation of Molecular Function GO terms enriched in genes whose expression is associated with ecological predictors in enterocytes 1. (D) REVIGO scatterplot representation of Biological Process GO terms enriched in genes whose expression is associated with ecological predictors in enterocytes 1. (E) Time-calibrated species tree with branches colored according to the mean relative rates of trait evolution for δ^15^N (left), and for expression of genes associated with δ^15^N (right). (F) Time-calibrated species tree with branches colored according to the mean relative rates of trait evolution for normalized gut length (left), and for expression of genes associated with normalized gut length (right). (G) Time-calibrated species tree with branches colored according to the mean relative rates of trait evolution for PC2 of LPJ morphology (left), and for expression of genes associated with PC2 of LPJ morphology (right). (H) Comparison of slopes (blue) of ecological space expansion over time for the six ecological predictors, between the observed data and the Brownian motion null model of trait evolution. Lineage accumulation through time derived from the species tree is shown in dark gray.

## Methods

### Experimental animals

A total of 31 fish specimens were selected for this study. The animals were kept in the fish facility of the Zoological Institute, University of Basel, Switzerland, holding animal experimentation permit 1010H, under a 12h-12h light-dark cycle at 25°C. All experimental procedures on these animals were performed using protocols approved by the Cantonal Veterinary office, Basel-Stadt, Switzerland (License number-3074/32560). Specific sample information (species, age and sex) can be found in Table S1.

### Fish dissection

All animals were systematically sampled in the morning, at least 12 hours after the last food intake, to optimize subsequent cleaning of the intestine. In brief, all individuals were sacrificed, followed by an incision in the spinal cord, in accordance with study permit 2317. The intestine of sacrificed animals was then dissected, placed in ice-cold phosphate buffered saline (PBS) and stored on ice until tissue dissociation.

### Generation of scRNAseq data

Individual intestines were carefully flushed with ice-cold PBS under binoculars, to remove all visible gut content, and cut into three equally long parts along the anterior-posterior axis, which were then dissociated individually. All intestinal samples (N = 75) were thoroughly chopped with a razor blade and dissociated at 25°C for 20 minutes with low agitation (100 rpm) in either 1 or 3 mL of pre-warmed DMEM/F-12 medium 1X (Fisher Scientific) supplemented with 1 mg / mL of Dispase II (Sigma). To ensure optimal dissociation of the tissue, samples were mixed by pipetting every 4 minutes throughout the enzymatic dissociation process. Suspensions were centrifuged at 500 rcf for 7 minutes at 4°C, resuspended in 10% Fetal Bovine Serum (FBS) in DMEM/F-12, filtered with a 20 µm cell strainer (CellTrics), centrifuged again with the same conditions, resuspended in 10% FBS in DMEM/F-12, and centrifuged one last time at low speed (200-250 rcf) for 5 minutes at 4°C. Depending on the size of each sample, the pellet was resuspended in ice-cold 100 - 1,000 µL 10% FBS in DMEM/F-12. Cell concentration and viability were estimated on a Cellometer K2 (Nexcelom). Cell viability was higher than 80% in all samples. For each intestine, the three independent gut samples were subsequently pooled, ensuring that each sample contributed equally to the final pool in number of cells.

All intestine cell suspensions were then diluted to a concentration of 500 - 1,500 cells / µL and loaded in a 10X Genomics Chromium for subsequent generation of single-cell cDNA (Chromium Next GEM Single Cell 3’ kit, 10X Genomics). Following the manufacturer’s instructions, dual index libraries were prepared and quantified either on a 2100 Bioanalyzer system (Agilent) or a Tapestation 4200 system (Agilent). Libraries were sequenced on a NovaSeq 6000 system (Illumina) at the Genomics Facility Basel, jointly operated by the University of Basel and the Department of Biosystems Science and Engineering (D-BSSE) of ETH Zürich. A total of 61 - 297 million reads (median 117 million reads) were sequenced for each library.

### Generation of bulk sequencing data

Six individuals from three cichlid species (*Astatotilapia burtoni, Eretmodus cyanostictus* and *Lepidiolamprologus attenuatus*) were sacrificed. Dissection of individuals and cleaning of the intestines were performed as described in the previous section. Each intestine was divided into five equally long parts along the anterior-posterior axis, and each section was processed independently. The sections were first mechanically chopped with a razor blade, then homogenized on a FastPrep 24 instrument (MP BioMedicals), and total RNA was extracted using a Zymo Direct-Zol RNA Miniprep kit (Zymo Research), following the manufacturer’s protocol. Illumina Truseq libraries (Illumina) were subsequently prepared and sequenced on a NovaSeq 6000 instrument (Illumina) at the joint Genomics Facility Basel, resulting in 11.4 - 17.3 million reads per sample.

### Raw scRNA data processing

The software Cell Ranger v5.0 (10X Genomics) was used to perform base calling, adaptor trimming and mapping the sequencing reads to the genome of the Nile tilapia (*O. niloticus*, GCA_001858045.3, RefSeq assembly GCF_001858045.2 (Conte *et al*., 2017)), a closely-related species evolutionarily equidistant to all Lake Tanganyika species included in this study, and for which a high-quality chromosome-level assembly and annotation are publicly available. 46.4% - 86.2% of sequencing reads (mean 67.5%) mapped confidently to the reference genome, and 32.5% - 75.6% (mean 52.8%) mapped confidently to the reference transcriptome (Table S2). We obtained a cell / gene count matrix for each sample, which consisted of 1,133-8,226 cells (mean 4,498 cells), with a means of 33,058 reads, 2,255 UMI counts, and 713 detected genes (Figure S1A-C). In total, we detected in each sample between 20,877 and 26,535 genes (mean 24,195 genes).

These cell / gene count matrices were filtered to eliminate low-quality cells. In short, we first corrected the gene expression data for ambient mRNA contamination based on the estimated mRNA expression profile from empty droplets, using the R package SoupX (Young & Behjati, 2020) (v1.6.2). We filtered out all cells with a mitochondrial contribution to UMI count higher than 80% and cells with a low ratio of detected genes per UMI counts. We followed the filtering procedure implemented in the R package ddqcR (Subramanian *et al*., 2022) (v0.1), which filters out cells with fewer than 100 detected genes, clusters the data into biologically meaningful clusters and performs a data-driven quality control at the level of clusters.

Subsequently, we performed normalization of the gene expression data from all retained cells of each sample and cell clustering using the R package Seurat (Satija *et al*., 2015) (v4.1.0). Specifically, we first used the SCTransform function(Hafemeister & Satija, 2019) to normalize the data, using the fraction of UMI counts mapping to mitochondrial genes, and the fraction of UMI counts mapping to ribosomal genes as regression variables. We then performed a Principal Component Analysis (PCA) using the RunPCA function of the R package Seurat, and the cells of each sample were clustered based on the first 30 components of the PCA using the FindNeighbors and FindClusters functions, with default parameters and a resolution of 1.

### Integration of single-cell datasets

All single-cell clustered and normalized datasets were integrated using the SelectIntegrationFeatures, PrepSCTIntegration, FindIntegrationAnchors, and IntegrateData functions from the R package Seurat (Satija *et al*., 2015; Stuart *et al*., 2019) (v4.1) with 5,000 features and a reference-based integration approach. The resulting integrated dataset was normalized and scaled, using the regression variables mentioned above and a cell cycle score computed using the CellCycleScoring function and a database of 87 Nile tilapia orthologs of zebrafish genes known to be involved in cell cycle. The resulting dataset was clustered based on the top 100 PCA components, based on the Louvain algorithm and with a resolution of 3, and visualized with the Uniform Manifold Approximation and Projection (UMAP) algorithm, using the RunUMAP function. We then filtered out non-informative clusters with a median combined mitochondrial and ribosomal fraction higher than 50%, performed an extra clustering step with similar parameters and obtained a final integrated dataset composed of 82,621 high-quality clustered cells derived from 25 species (Figure 1B).

### Identification of biological cell types

Following the initial clustering, we identified all major clusters as cell type categories (Figure 1B,C). Cell population characterizations were performed within each cell type category, using a finer sub-clustering with low resolution parameters (0.1-0.4). Cell types were identified based on the expression of cell type marker genes described in the literature, and by contrasting uncorrected gene expression patterns between clusters, as performed in the FindAllMarkers function from the R package Seurat, with the following parameters: min.pct = 0.25, logfc.threshold = 0.5, method = “MAST” (Figure 1C). Subsequently, cross-species identification of cell types was validated using functions from the R packages SciBet (Li *et al*., 2020) (v1.0) and MetaNeighbor (Crow *et al*., 2018) (v1.14) (Figure S1E). In brief, the MetaNeighborUS function was used to compute cell type to cell type AUROC scores and verify that identified cell types were robust to cross-species differences. The topHits function was then called with a threshold parameter of 0.8 to visually inspect for potential high matches between different cell types. The conf function from the R package SciBet was used to generate a confusion matrix that evaluates the performance of the cell type classification, to ensure that diagonal values, which correspond to self-validation scores, were higher than 0.8. To further characterize the identified cell types, we estimated the fraction of reads mapping to mitochondrial genes, and counted the number of counts (UMIs) and detected genes in each cell of each cell type (Figure S1A-C). Finally, for each cell type, we calculated the proportions of transcripts that show high cell-population specificity in expression, as defined by a gsi higher than 6. This revealed a variable number of cell-population specific features among cell types (Figure S1D).

### Characterization of epithelial cells

We identified four main populations of enterocytes in our single-cell datasets, which we further characterized by examining the expression of 1-to-1 orthologs of zebrafish genes that show conserved regional transcriptional specification along the intestine in zebrafish and mouse (Figure S2A). This analysis revealed strong anterior-posterior signatures for enterocytes 1, and to a lesser extent, in enterocytes 2 and lysosome-rich enterocytes.

To confirm these findings, we applied spatially defined bulk mRNA sequencing from three species included in the single-cell dataset, see section ‘Generation of bulk sequencing data’ in Method details. The obtained sequencing reads were trimmed using Trimmomatic (Bolger, Lohse & Usadel, 2014) (v0.39), and mapped to the genome of the Nile tilapia using STAR (Dobin *et al*., 2013) (v2.7.3) and HTSeq (Putri *et al*., 2022) (v2.0) (Table S3). We then used the R package BayesPrism (Chu *et al*., 2022) (v2.0) to deconvolute the bulk in each part of each species separately, and predict cellular composition in cell types from total RNA sequencing data, using both mRNA and single-cell raw counts as input (Figure S2B,C). This revealed strong anterior-posterior identity in enterocytes 1, enterocytes 2 and lysosome-rich enterocytes.

To characterize the cellular and molecular functions inherent to the three major types of enterocytes, we calculated pseudobulk values for each gene in each cell type, and contrasted the gene expression signatures through differential expression analysis. Finally, we performed gene ontology (GO)(Gene Ontology Consortium *et al*., 2023) term enrichment analyses on the sets of differentially expressed genes using the R packages limma (Ritchie *et al*., 2015) (v3.50.3) and GO.db (v3.14), and visualized these using the REVIGO web server (Supek *et al*., 2011). This revealed the molecular functions, cellular components and biological processes associated with each type of enterocytes and by extension, with each part of the intestine (Figure S2D-G).

### Characterization of gene expression signatures

We first assessed gene expression affinities between cell populations by calculating Pearson pairwise correlations between cell populations and a hierarchical cluster analysis based on the ward D2 agglomeration method (Figure 1D). In addition, for each cell type and species, we generated pseudobulk samples by aggregating raw counts across cells, using the GetAssayData function from the R package Seurat. We filtered out from the intestine matrix of pseudobulk samples lowly expressed genes that are not expressed in at least two cells of any cell type, across all species. The resulting data consisted of 13,150 genes, whose expression levels were quantified in all 24 identified cell populations. To visualize global patterns of gene expression differences across species and cell populations, the gene expression data was further normalized using the median of ratios method implemented in the R package DESeq2 (Love, Huber & Anders, 2014) (v1.14) to account for between-sample differences in sequencing depth and RNA composition. We next transformed the data using the variance-stabilizing transformation method as implemented by the vst function from DESeq2 and performed a PCA based on the top 10,000 variable genes using the prcomp function (Figure 1E, Figure S2C,D).

Subsequently, the pseudobulk expression values were normalized to counts per million (cpm) values, which were used for all downstream analyses. To account for potential differences and biases in expression quantification across species and single-cell data generation batches, we calculated a gene specificity index for each gene, species and cell type, as defined by Tosches and colleagues (Tosches *et al*., 2018); Figure S1F). In short, for every species included, we computed for each gene the ratio between its level expression within each cell type and its mean expression across all cell types, and performed pairwise correlations across all species and cell types, using Spearman’s rank correlation coefficients.

### Cross-species gene expression divergence

We calculated cross-species gene expression in a pairwise fashion and within each cell population, after filtering out pseudobulks aggregated on fewer than 20 cells, and cell populations with fewer than 17 species represented by at least 20 cells. This resulted in a dataset consisting of cpm values calculated in 18 cell populations, with varying numbers of species represented in these cell populations. For each pair of species and within each cell population, we computed the mean-squared distances as implemented in EVEE-tools (Chen *et al*., 2019); Figure 2A,B). To investigate whether gene expression divergence tends to correlate positively with phylogenetic divergence, we also retrieved published evolutionary distances between all pairs of species in the dataset, calculated as genomic substitutions across the genome (Ronco *et al*., 2021) and plotted gene expression mean squared distances as a function of evolutionary distances, revealing a non-linear, positive correlation between them (Figure S3).

Cross-species gene expression divergence was also estimated with the R package Augur (Skinnider *et al*., 2021) (v1.0.3), using a method designed for prioritizing cell type responses to experimental conditions, with conditions set to ‘species’ here. In short, Augur tries to predict what species each cell from a given cell type comes from, based on gene expression, and gives an overall AUC (Area Under Curve), a confidence score that ranges from 0.5, if the species identity cannot be recovered better than randomly, to 1, if the species identity is always correctly guessed.

To account for potential differences in sequencing depth, each cell was downsampled to the same number of UMIs (1000 UMIs per cell). The Augur method was applied to the integrated dataset restricted to the 18 selected cell populations with enough species represented by 20 cells or more. Augur was then run in a pairwise fashion, in subsets of the dataset containing only two of the species, with the following parameters: subsample_size = 10, min_cells = 10, n_subsamples = 50. Pairwise AUC scores were compared across the 18 selected cell populations, as well as across pairs of species and phylogenetic distances separating these pairs (Figure 2C).

### Cell type-specific transcriptomes

We used four different approaches to characterize the genes expressed in each cell population, and the evolutionary forces at play that may constrain the evolution of cell populations and of their associated transcriptomes. First, we reconstructed the transcriptome age index (TAI) of each gene in the genome of the Nile tilapia. This is a measure of the evolutionary age, whereby higher TAI values indicate younger genes and lower TAI values the older ones. To do so, we followed the procedure implemented in the Python package oggmap (Ullrich & Glytnasi, 2023), which relies on a phylostratigraphic approach to extract gene ages in orthology groups, enabling us to calculate the evolutionary ages for 12,550 such orthologs expressed in our intestine single-cell dataset (Figure 2D). Second, we computed tissue-specificity indices (Tau, *τ*), a measure of how tissue-specific the expression of a gene, which ranges from 0 to 1. We used the procedure from El Taher and colleagues (El Taher *et al*., 2021), based on gene expression data derived from 13 tissues of cichlid fishes: brain, retina, gonads, lower pharyngeal jaw, gills, liver, intestine, skin, spleen, kidney, heart, muscle and blood (Brawand *et al*., 2011; Xia *et al*., 2018; Xu *et al*., 2018; Ellison *et al*., 2018; Rajkov *et al*., 2021; El Taher *et al*., 2021; Ricci *et al*., 2023). This resulted in *τ* values for 25,687 genes expressed in the intestine (Figure 2E).

Third, we retrieved predicted protein-protein interactions (PPIs) in *O. niloticus* from the STRING v11 database (Szklarczyk *et al*., 2022), and selected high-confidence PPIs, as indicated by a score higher than 0.7 (Bozhilova *et al*., 2019). This resulted in high-confidence PPI predictions for a total of 13,977 PCGs expressed in the intestine (Figure 2F).

Fourth, we retrieved from Roux and Robinson-Rechavi(Roux & Robinson-Rechavi, 2008) ratios of non-synonymous to synonymous substitution (dN/dS), calculated for 4172 orthologous protein-coding gene among three species of distantly-related teleost fish species, namely *Tetraodon nigroviridis*, *Takifugu rubripes* and *Danio rerio*. We used dN/dS as a proxy for the evolutionary selective pressure acting on protein-coding genes. For each cell present in the integrated dataset, we weighted each individual gene score (age score, by its expression value within the cell and computed the overall cell transcriptome age index, tissue specificity score and dN/dS by calculating the mean of these weighted scores across all genes expressed (i.e. with UMI count > 1; Figure 2G).

Finally, to summarize these potential constraints acting on the transcriptomes of intestinal cell populations and the cross-species divergences, we performed a PCA using all six traits described above, namely mean MSGED, mean AUC, mean TAI, mean Tau, mean PPI and mean dN/dS (Figure 2H).

### Detection of gene co-expression modules

We detected gene co-expression modules using the R package scWGCNA (Feregrino & Tschopp, 2022), an adaptation of WGCNA (Langfelder & Horvath, 2008) for single-cell datasets. In total, we identified 25 modules of co-expressed genes, comprising 17 - 103 genes each (mean = 53 genes; Figure S1I). We next quantified module expression in each identified cell type to characterize the cell type specificity and molecular functions of each scWGCNA module. GO term enrichment analyses were then performed using the R packages limma and GO.db, and visualized using the REVIGO web server (Supek *et al*., 2011), to highlight molecular functions associated with the modules of co-expressed genes (Figure S1J,K).

### Ecological predictors

We used a total of six eco-morphological proxies. First, we retrieved the carbon (δ^13^C-values) and nitrogen (δ^15^N-values) stable isotope compositions of muscle tissue, previously measured for all cichlid species of the Lake Tanganyika radiation (Ronco *et al*., 2021), as proxies for their macroecology, namely the position along the benthic-pelagic axis, and trophic level, respectively. Subsequently, we retrieved gut length data across 126 species from Duque-Correa et al. 2024 (Duque-Correa *et al*., 2024), and calculated the normalized gut length (so called Zihler index (Zihler, 1981), Figure S4A), and included two other morphological predictors in our analyses: PC2 of lower pharyngeal jaw morphology and PC1 of oral jaw morphology, measured across virtually all species of the Lake Tanganyika adaptive radiation (Ronco *et al*., 2021), which have been shown to be mostly associated with trophic ecology and food uptake, respectively. Finally, we reconstructed a diet ecology vector, based on a PCA calculated from nitrogen stable isotope data, lower pharyngeal jaw morphology and normalized gut length data (Figure S4B,C).

### Ecological signals in cell type abundances

For each specimen, we estimated the relative abundances of the different cell populations using the plotCellTypeProps function from the R package speckle (Phipson *et al*., 2022) (v0.0.3). Because the efficiency of dissociation of epithelium and lamina propria can vary from species to species, due to biases in intestine thickness, robustness and elasticity, we focused on the epithelial cell types, which form a single layer of cells, separating the intestinal lumen from the lamina propria. Specifically, we calculated the relative abundance of all epithelial absorptive and non-absorptive cells within the epithelium (Figure 3A). To test for association between relative cell type abundances and ecological predictors, we applied a linear regression (lm function in R), and a phylogenetic generalized least-square analysis (pgls function from the R package caper (Orme, 2023) (v1.0.3)), which accounts for the phylogenetic non-independence of the species. We then reported the coefficients of determination and statistical p-values to identify cell types and cell categories whose differences in relative abundances are ecologically relevant (Figure 3B-D).

### Associations between ecology and cell population

To investigate associations between gene expression in intestinal cell populations and ecological predictors, we created three sets of genes. For each cell population, we first selected all genes expressed in at least 50% of the species and in 20% of the cells, which will be referred to as the ‘global’ gene set. In parallel, we established a ‘specific’ gene set, consisting only of cell-type specific genes (as defined in section ‘Characterization of cell population-specific transcriptomes’. Finally, we constructed a third ‘module’ gene set, consisting of the genes of scWGCNA modules only. We then used the cpm pseudobulk dataset presented in section ‘Cross-species cell population gene expression divergence’, consisting of 18 cell populations and with varying number of species per cell population, according to how many of these species are represented by at least 20 cells and by a sufficient number of counts in each cell population. This gene set was log10-transformed and filtered for each one of the three sets of genes, resulting in a ‘global’ gene set consisting of 18 cell populations and 454 – 3,221 genes (mean = 1607 genes) per cell population and a ‘specific’ gene set, consisting of 18 cell populations and 30 – 360 genes (mean = 132 genes) per cell population. To build the ‘module’ gene set, for each module we selected the cell population in which the module expression is the highest, and filtered out 3 modules with either low cell population specificity or high specificity in a very rare cell population. This resulted in a ‘module’ gene set consisting of 22 gene expression matrices with 17 – 103 genes per matrix (mean = 53 genes).

For each one of the three gene sets, we assessed the association between each ecological predictor and cell population gene expression using a multivariate phylogenetic regression approach, as implemented in the functions mvgls and manova.gls of the R package mvMORPH (Clavel *et al*., 2015; Clavel, Aristide & Morlon, 2019; Clavel & Morlon, 2020) (v1.1.8). Specifically, we fitted a Brownian motion (BM) model of evolution with a Pagel’s lambda phylogenetic tree transformation and a penalized likelihood approach, and applied a type II phylogenetic MANOVA using the Pillai’s test statistic and 1000 permutations to detect potential significant correlations (Figure 4A-C).

To identify which ecological predictors are most relevant in ecology-gene expression correlations, we performed phylogenetic two-block partial least-squares (p2B-PLS) analysis, as implemented in the two.b.pls function of the R package geomorph (Baken *et al*., 2021) (v.4.0.7). The multivariate predictor matrix consisted in the nitrogen and carbon stable isotopes, as well as gut normalized gut length, PC2 of lower pharyngeal jaw morphology and PC1 of oral jaw morphology. The individual ecological predictor loadings were then compared in each cell population of each one of the three datasets (Figure S4D-F).

### Characterizing ecologically relevant genes

To characterize associations between ecological predictors and gene expression profiles, we investigated correlations at the individual gene level in ecologically relevant cell populations. Within each epithelial cell population, we conducted linear regression (lm) and pGLS analyses between each individual ecological predictor and all individual genes expressed in at least 10% of cells and across at least 50% of species. Specifically, we tested for correlations between the predictor values and the log10-transformed cpm pseudobulk values, using the time-calibrated genome-wide species tree of Lake Tanganyika cichlid species to account for the non-phylogenetic independence of the species. For each ecological predictor, we selected the genes showing significant correlations in both lm and pGLS analyses (p-values < 0.05, Figure S5A).

Additionally, we performed a two-block partial least-squares (2B-PLS) analysis, as well as a phylogenetic two-block partial least-squares (p2B-PLS) analysis, using as input each ecological predictor individually and the overall log-transformed filtered cpm pseudobulk gene expression matrix for the cell population enterocytes 1, i.e. the cell population showing the most significant associations with dietary predictors, using the 2b.pls and phylo.integrations functions from the R package geomorph (Baken *et al*., 2021). We then identified the top genes driving the correlations between the two blocks, selecting the same number of genes as those correlating with predictors at the individual levels in lm and pGLS analyses, and calculated the intersection between (i) the significant genes in lm and pGLS analyses, (ii) the top 2B-PLS genes, and (iii) the top p2B-PLS genes, to create sets of genes correlating with ecological predictors both at the univariate and multivariate levels, and both with and without accounting for the phylogenetic non-independence of the samples during data rotation (Figure S5B).

We further characterized the ecologically relevant genes in enterocytes 1 by comparing them to all genes expressed in at least 10% of cells in each cell population in terms of tissue specificity (Tau; Figure 5A) and gene specificity index (gsi; Figure 5B). We performed GO enrichment analysis to identify molecular functions and biological processes enriched in genes whose expression correlated positively and negatively with trophic levels, respectively. Finally, we visualized these using the REVIGO web server (Supek *et al*., 2011) (Figure S5C,D).

### Reconstruction of gene expression and trait evolution

For each cell population and expressed gene, we modelled the evolution of gene expression across the species phylogeny, as defined by the time-calibrated genome-wide phylogenetic tree (Ronco *et al*., 2021). To do so, we first excluded the outgroup Nile tilapia, and genes expressed in fewer than 50% of the species and in less than 10% of cells. We log-transformed the filtered cpm pseudobulk gene expression data and used the fitContinuous function from the R package geiger(Harmon *et al*., 2008) (v2.0.7) to fit a Brownian motion (BM) mode of evolution on each expressed gene individually. A rate of gene expression evolution was inferred from the output of fitContinuous for each expressed gene in enterocytes 1, as well as in the ecologically relevant genes (Figure 5C).

To reconstruct the evolution of entire sets of ecologically-relevant genes along the phylogeny of Lake Tanganyika cichlid fishes and identify lineage-specific rates of evolution, we first calculated the median value of gene expression for each set of genes (Figure 5D), and mapped the evolution of the median gene expression values onto the time-calibrated phylogenetic species tree (Figure 5E,F) by modeling gene expression evolution as a Brownian motion process and estimating internal node values using a maximum-likelihood (ML) approach, as implemented in the contMap function from the R package phytools(Revell, 2024) (v2.1.1). We then visualized the rates of gene expression evolution using a custom R script (Figure 5F, Figure S5E-G).

Finally, we estimated branch-specific rates for each ecological predictor – namely δ^15^N, δ^13^C, normalized gut length, PC2 of LPJ morphology, PC1 of oral jaw morphology, and the diet ecology vector – by fitting a variable rates model using BayesTraits (Venditti, Meade & Pagel, 2011) (v3.0.2). For each trait we ran a single MCMC chain for at least 1,000,000,000 iterations, sampling parameters every 100,000 iterations after discarding the first 10,000,000 iterations as a burn-in. Stationary of each chain was accepted when the effective sample size (ESS) of each parameter reached at least 200 (calculated using the R package coda (Plummer *et al*., 2006) (v0.19-3). For the parameters alpha (ancestral value) and the rate parameter sigma, we defined uniform prior distributions per trait (δ^13^C: alpha = -50-0; sigma = 0-10; δ^15^N: alpha = 0-20; sigma = 0-10; normalized gut length: alpha = -10-5; sigma = 0-20; diet ecology vector: alpha = -10-10; sigma = 0-10; LPJ PC2 and oral jaw PC1 = alpha = -1-1; sigma = 0-0.001) that were informed from a single process Brownian motion model, fitted using the R package geiger. For each ecological predictor, we then extracted branch-specific evolutionary rates from the mean rate-transformed tree of the posterior sample and reconstructed ML ancestral states using the R package phytools and compared it to the tree of median expression of ecologically-relevant genes using a linear model (Figure 5F, Figure S5E-G). To compare the diversity accumulation over time relative to a null model of trait evolution (Cooney *et al*., 2017; Ronco *et al*., 2021), we further reconstructed rate-informed maximum likelihood ancestral states using the R package phytools and calculated ecological diversity in time intervals of 0.15 million years, once from the ancestral states of the observed data and once based on ancestral states from 500 datasets that were simulated under Brownian motion. Finally, we compared the slopes of the diversity accumulation over time of observed data with each of the null models (observed – BM; Figure S5H).

## References

Abzhanov, A., Kuo, W.P., Hartmann, C., Grant, B.R., Grant, P.R. & Tabin, C.J. (2006) The calmodulin pathway and evolution of elongated beak morphology in Darwin’s finches. Nature 442, 563–567.

Abzhanov, A., Protas, M., Grant, B.R., Grant, P.R. & Tabin, C.J. (2004) Bmp4 and morphological variation of beaks in Darwin’s finches. Science 305, 1462–1465.

Baken, E.K., Collyer, M.L., Kaliontzopoulou, A. & Adams, D.C. (2021) geomorph v4.0 and gmShiny: Enhanced analytics and a new graphical interface for a comprehensive morphometric experience. Methods in ecology and evolution / British Ecological Society 12, 2355–2363.

Bakke-McKellep, A.M., Penn, M.H., Salas, P.M., Refstie, S., Sperstad, S., Landsverk, T., Ringø, E. & Krogdahl, A. (2007) Effects of dietary soyabean meal, inulin and oxytetracycline on intestinal microbiota and epithelial cell stress, apoptosis and proliferation in the teleost Atlantic salmon (Salmo salar L.). The British journal of nutrition 97, 699–713.

Baldo, L., Pretus, J.L., Riera, J.L., Musilova, Z., Bitja Nyom, A.R. & Salzburger, W. (2017) Convergence of gut microbiotas in the adaptive radiations of African cichlid fishes. The ISME journal 11, 1975–1987.

Bedford, T. & Hartl, D.L. (2009) Optimization of gene expression by natural selection. Proceedings of the National Academy of Sciences of the United States of America 106, 1133–1138.

Berthelot, C., Villar, D., Horvath, J.E., Odom, D.T. & Flicek, P. (2018) Complexity and conservation of regulatory landscapes underlie evolutionary resilience of mammalian gene expression. Nature ecology & evolution 2, 152–163.

Bolger, A.M., Lohse, M. & Usadel, B. (2014) Trimmomatic: a flexible trimmer for Illumina sequence data. Bioinformatics 30, 2114–2120.

Bozhilova, L.V., Whitmore, A.V., Wray, J., Reinert, G. & Deane, C.M. (2019) Measuring rank robustness in scored protein interaction networks. BMC bioinformatics 20, 446.

Brawand, D., Soumillon, M., Necsulea, A., Julien, P., Csárdi, G., Harrigan, P., Weier, M., Liechti, A., Aximu-Petri, A., Kircher, M., Albert, F.W., Zeller, U., Khaitovich, P., Grützner, F., Bergmann, S., et al. (2011) The evolution of gene expression levels in mammalian organs. Nature 478, 343–348.

Brawand, D., Wagner, C.E., Li, Y.I., Malinsky, M., Keller, I., Fan, S., Simakov, O., Ng, A.Y., Lim, Z.W., Bezault, E., Turner-Maier, J., Johnson, J., Alcazar, R., Noh, H.J., Russell, P., et al. (2014) The genomic substrate for adaptive radiation in African cichlid fish. Nature 513, 375–381.

Calduch-Giner, J.A., Sitjà-Bobadilla, A. & Pérez-Sánchez, J. (2016) Gene expression profiling reveals functional specialization along the intestinal tract of a carnivorous teleostean fish (*Dicentrarchus labrax)*. Frontiers in physiology 7, 359.

Cardoso-Moreira, M., Halbert, J., Valloton, D., Velten, B., Chen, C., Shao, Y., Liechti, A., Ascenção, K., Rummel, C., Ovchinnikova, S., Mazin, P.V., Xenarios, I., Harshman, K., Mort, M., Cooper, D.N., et al. (2019) Gene expression across mammalian organ development. Nature 571, 505–509.

Carroll, S.B. (2005) Evolution at two levels: on genes and form. PLoS biology 3, e245.

Chen, J., Swofford, R., Johnson, J., Cummings, B.B., Rogel, N., Lindblad-Toh, K., Haerty, W., Palma, F. di & Regev, A. (2019) A quantitative framework for characterizing the evolutionary history of mammalian gene expression. Genome research 29, 53–63.

Childers, L., Park, E., Wang, S., Liu, R., Barry, R., Watts, S.A., Rawls, J.F. & Bagnat, M. (2024) Protein absorption in the zebrafish gut is regulated by interactions between lysosome rich enterocytes and the microbiome. bioRxiv.

Chu, T., Wang, Z., Pe’er, D. & Danko, C.G. (2022) Cell type and gene expression deconvolution with BayesPrism enables Bayesian integrative analysis across bulk and single-cell RNA sequencing in oncology. Nature cancer 3, 505–517.

Clara, R., Schumacher, M., Ramachandran, D., Fedele, S., Krieger, J.-P., Langhans, W. & Mansouri, A. (2017) Metabolic adaptation of the small intestine to short- and medium-term high-fat diet exposure. Journal of cellular physiology 232, 167–175.

Clavel, J., Aristide, L. & Morlon, H. (2019) A penalized likelihood framework for high-dimensional phylogenetic comparative methods and an application to New-World monkeys brain evolution. Systematic biology 68, 93–116.

Clavel, J., Escarguel, G. & Merceron, G. (2015) Mvmorph: An r package for fitting multivariate evolutionary models to morphometric data. Methods in ecology and evolution / British Ecological Society 6, 1311–1319.

Clavel, J. & Morlon, H. (2020) Reliable phylogenetic regressions for multivariate comparative data: illustration with the manova and application to the effect of diet on mandible morphology in Phyllostomid bats. Systematic biology 69, 927–943.

Conte, M.A., Gammerdinger, W.J., Bartie, K.L., Penman, D.J. & Kocher, T.D. (2017) A high quality assembly of the Nile Tilapia (*Oreochromis niloticus*) genome reveals the structure of two sex determination regions. BMC genomics 18, 341.

Cooney, C.R., Bright, J.A., Capp, E.J.R., Chira, A.M., Hughes, E.C., Moody, C.J.A., Nouri, L.O., Varley, Z.K. & Thomas, G.H. (2017) Mega-evolutionary dynamics of the adaptive radiation of birds. Nature 542, 344–347.

Crosnier, C., Vargesson, N., Gschmeissner, S., Ariza-McNaughton, L., Morrison, A. & Lewis, J. (2005) Delta-Notch signalling controls commitment to a secretory fate in the zebrafish intestine. Development 132, 1093–1104.

Cross, F.R., Buchler, N.E. & Skotheim, J.M. (2011) Evolution of networks and sequences in eukaryotic cell cycle control. Philosophical transactions of the Royal Society of London. Series B, Biological sciences 366, 3532–3544.

Crow, M., Paul, A., Ballouz, S., Huang, Z.J. & Gillis, J. (2018) Characterizing the replicability of cell types defined by single cell RNA-sequencing data using MetaNeighbor. Nature communications 9, 884.

Darwin, C. (1859) On The Origin Of Species. London: John Murray.

Dobin, A., Davis, C.A., Schlesinger, F., Drenkow, J., Zaleski, C., Jha, S., Batut, P., Chaisson, M. & Gingeras, T.R. (2013) STAR: ultrafast universal RNA-seq aligner. Bioinformatics 29, 15–21.

Domazet-Lošo, T. & Tautz, D. (2010) A phylogenetically based transcriptome age index mirrors ontogenetic divergence patterns. Nature 468, 815–818.

Duque-Correa, M.J., Clements, K.D., Meloro, C., Ronco, F., Boila, A., Indermaur, A., Salzburger, W. & Clauss, M. (2024) Diet and habitat as determinants of intestine length in fishes. Reviews in fish biology and fisheries 34, 1017–1034.

Ellison, A.R., Uren Webster, T.M., Rey, O., Garcia de Leaniz, C., Consuegra, S., Orozco-terWengel, P. & Cable, J. (2018) Transcriptomic response to parasite infection in Nile tilapia (*Oreochromis niloticus*) depends on rearing density. BMC genomics 19, 723.

Elmentaite, R., Kumasaka, N., Roberts, K., Fleming, A., Dann, E., King, H.W., Kleshchevnikov, V., Dabrowska, M., Pritchard, S., Bolt, L., Vieira, S.F., Mamanova, L., Huang, N., Perrone, F., Goh Kai’En, I., et al. (2021) Cells of the human intestinal tract mapped across space and time. Nature 597, 250–255.

El Taher, A., Böhne, A., Boileau, N., Ronco, F., Indermaur, A., Widmer, L. & Salzburger, W. (2021) Gene expression dynamics during rapid organismal diversification in African cichlid fishes. Nature ecology & evolution 5, 243–250.

Engel, J., Blanchet, L., Bloemen, B., van den Heuvel, L.P., Engelke, U.H.F., Wevers, R.A. & Buydens, L.M.C. (2015) Regularized MANOVA (rMANOVA) in untargeted metabolomics. Analytica chimica acta 899, 1–12.

Enriquez, J.R., McCauley, H.A., Zhang, K.X., Sanchez, J.G., Kalin, G.T., Lang, R.A. & Wells, J.M. (2022) A dietary change to a high-fat diet initiates a rapid adaptation of the intestine. Cell reports 41, 111641.

Farache, J., Koren, I., Milo, I., Gurevich, I., Kim, K.-W., Zigmond, E., Furtado, G.C., Lira, S.A. & Shakhar, G. (2013) Luminal bacteria recruit CD103+ dendritic cells into the intestinal epithelium to sample bacterial antigens for presentation. Immunity 38, 581–595.

Fay, J.C. & Wittkopp, P.J. (2008) Evaluating the role of natural selection in the evolution of gene regulation. Heredity 100, 191–199.

Feregrino, C. & Tschopp, P. (2022) Assessing evolutionary and developmental transcriptome dynamics in homologous cell types. Developmental dynamics 251, 1472–1489.

Fischer, M., Schade, A.E., Branigan, T.B., Müller, G.A. & DeCaprio, J.A. (2022) Coordinating gene expression during the cell cycle. Trends in biochemical sciences 47, 1009–1022.

Fre, S., Bardin, A., Robine, S. & Louvard, D. (2011) Notch signaling in intestinal homeostasis across species: the cases of Drosophila, Zebrafish and the mouse. Experimental cell research 317, 2740–2747.

Fryer, G. & Iles, T.D. (1972) The Cichlid Fishes Of The Great Lakes Of Africa. T.F.H. Publications.

Futuyma, D.J. & Moreno, G. (1988) The evolution of ecological specialization. Annual review of ecology and systematics 19, 207–233.

Gavrilets, S. & Losos, J.B. (2009) Adaptive radiation: contrasting theory with data. Science 323, 732–737.

Gene Ontology Consortium, Aleksander, S.A., Balhoff, J., Carbon, S., Cherry, J.M., Drabkin, H.J., Ebert, D., Feuermann, M., Gaudet, P., Harris, N.L., Hill, D.P., Lee, R., Mi, H., Moxon, S., Mungall, C.J., et al. (2023) The Gene Ontology knowledgebase in 2023. Genetics 224, iyad031.

Grant, P.R. (1986) Ecology And Evolution Of Darwin’s Finches. Princeton University Press, Princeton, NJ, USA

Grant, P.R. & Grant, B.R. (2006) Evolution of character displacement in Darwin’s finches. Science 313, 224–226.

Greenwood-Van Meerveld, B., Johnson, A.C. & Grundy, D. (2017) Gastrointestinal physiology and function. Handbook of experimental pharmacology 239, 1–16.

Haber, A.L., Biton, M., Rogel, N., Herbst, R.H., Shekhar, K., Smillie, C., Burgin, G., Delorey, T.M., Howitt, M.R., Katz, Y., Tirosh, I., Beyaz, S., Dionne, D., Zhang, M., Raychowdhury, R., et al. (2017) A single-cell survey of the small intestinal epithelium. Nature 551, 333–339.

Hafemeister, C. & Satija, R. (2019) Normalization and variance stabilization of single-cell RNA-seq data using regularized negative binomial regression. Genome biology 20, 296.

Hansen, T.F. (1997) Stabilizing selection and the comparative analysis of adaptation. Evolution; international journal of organic evolution 51, 1341–1351.

Harmon, L.J., Weir, J.T., Brock, C.D., Glor, R.E. & Challenger, W. (2008) GEIGER: investigating evolutionary radiations. Bioinformatics 24, 129–131.

Harper, J., Mould, A., Andrews, R.M., Bikoff, E.K. & Robertson, E.J. (2011) The transcriptional repressor Blimp1/Prdm1 regulates postnatal reprogramming of intestinal enterocytes. Proceedings of the National Academy of Sciences of the United States of America 108, 10585–10590.

He, S., Liang, X.-F., Li, L., Sun, J., Wen, Z.-Y., Cheng, X.-Y., Li, A.-X., Cai, W.-J., He, Y.-H., Wang, Y.-P., Tao, Y.-X. & Yuan, X.-C. (2015) Transcriptome analysis of food habit transition from carnivory to herbivory in a typical vertebrate herbivore, grass carp *Ctenopharyngodon idella*. BMC genomics 16, 15.

Hill, M.S., Vande Zande, P. & Wittkopp, P.J. (2021) Molecular and evolutionary processes generating variation in gene expression. Nature reviews. Genetics 22, 203–215.

Hino, S., Takemura, N., Sonoyama, K., Morita, A., Kawagishi, H., Aoe, S. & Morita, T. (2012) Small intestinal goblet cell proliferation induced by ingestion of soluble and insoluble dietary fiber is characterized by an increase in sialylated mucins in rats. The Journal of nutrition 142, 1429–1436.

Hodgkin, J. (1998) Seven types of pleiotropy. The International journal of developmental biology 42, 501–505.

Hopperdietzel, C., Hirschberg, R.M., Hünigen, H., Wolter, J., Richardson, K. & Plendl, J. (2014) Gross morphology and histology of the alimentary tract of the convict cichlid *Amatitlania nigrofasciata*. Journal of fish biology 85, 1707–1725.

Huang, X., Zhong, L., Kang, Q., Liu, S., Feng, Y., Geng, Y., Chen, D., Ou, Y., Yang, S., Yin, L. & Luo, W. (2021) A high starch diet alters the composition of the intestinal microbiota of largemouth bass *Micropterus salmoides*, which may be associated with the development of enteritis. Frontiers in microbiology 12, 696588.

Hung, R.-J., Hu, Y., Kirchner, R., Liu, Y., Xu, C., Comjean, A., Tattikota, S.G., Li, F., Song, W., Ho Sui, S. & Perrimon, N. (2020) A cell atlas of the adult Drosophila midgut. Proceedings of the National Academy of Sciences of the United States of America 117, 1514–1523.

Hunnicutt, K.E., Good, J.M. & Larson, E.L. (2022) Unraveling patterns of disrupted gene expression across a complex tissue. Evolution; international journal of organic evolution 76, 275–291.

Ito, H., Satsukawa, M., Arai, E., Sugiyama, K., Sonoyama, K., Kiriyama, S. & Morita, T. (2009) Soluble fiber viscosity affects both goblet cell number and small intestine mucin secretion in rats. The Journal of nutrition 139, 1640–1647.

Johnson, Z.V., Hegarty, B.E., Gruenhagen, G.W., Lancaster, T.J., McGrath, P.T. & Streelman, J.T. (2023) Cellular profiling of a recently-evolved social behavior in cichlid fishes. Nature communications 14, 4891.

Jones, L.O., Willms, R.J., Xu, X., Graham, R.D.V., Eklund, M., Shin, M. & Foley, E. (2023) Single-cell resolution of the adult zebrafish intestine under conventional conditions and in response to an acute *Vibrio cholerae* infection. Cell reports 42, 113407.

Karasov, W.H., Martínez del Rio, C. & Caviedes-Vidal, E. (2011) Ecological physiology of diet and digestive systems. Annual review of physiology 73, 69–93.

Lamichhaney, S., Berglund, J., Almén, M.S., Maqbool, K., Grabherr, M., Martinez-Barrio, A., Promerová, M., Rubin, C.-J., Wang, C., Zamani, N., Grant, B.R., Grant, P.R., Webster, M.T. & Andersson, L. (2015) Evolution of Darwin’s finches and their beaks revealed by genome sequencing. Nature 518, 371–375.

Langfelder, P. & Horvath, S. (2008) WGCNA: an R package for weighted correlation network analysis. BMC bioinformatics 9, 559.

Lickwar, C.R., Camp, J.G., Weiser, M., Cocchiaro, J.L., Kingsley, D.M., Furey, T.S., Sheikh, S.Z. & Rawls, J.F. (2017) Genomic dissection of conserved transcriptional regulation in intestinal epithelial cells. PLoS biology 15, e2002054.

Li, C., Liu, B., Kang, B., Liu, Z., Liu, Y., Chen, C., Ren, X. & Zhang, Z. (2020) SciBet as a portable and fast single cell type identifier. Nature communications 11, 1818.

Liem, K.F. (1973) Evolutionary strategies and morphological innovations: cichlid pharyngeal jaws. Systematic zoology 22, 425.

Løkka, G. & Koppang, E.O. (2016) Antigen sampling in the fish intestine. Developmental and comparative immunology 64, 138–149.

Love, M.I., Huber, W. & Anders, S. (2014) Moderated estimation of fold change and dispersion for RNA-seq data with DESeq2. Genome biology 15, 550.

Luca, F., Perry, G.H. & Di Rienzo, A. (2010) Evolutionary adaptations to dietary changes. Annual review of nutrition 30, 291–314.

Mallarino, R., Grant, P.R., Grant, B.R., Herrel, A., Kuo, W.P. & Abzhanov, A. (2011) Two developmental modules establish 3D beak-shape variation in Darwin’s finches. Proceedings of the National Academy of Sciences of the United States of America 108, 4057–4062.

Muncan, V., Heijmans, J., Krasinski, S.D., Büller, N.V., Wildenberg, M.E., Meisner, S., Radonjic, M., Stapleton, K.A., Lamers, W.H., Biemond, I., van den Bergh Weerman, M.A., O’Carroll, D., Hardwick, J.C., Hommes, D.W. & van den Brink, G.R. (2011) Blimp1 regulates the transition of neonatal to adult intestinal epithelium. Nature communications 2, 452.

Murat, F., Mbengue, N., Winge, S.B., Trefzer, T., Leushkin, E., Sepp, M., Cardoso-Moreira, M., Schmidt, J., Schneider, C., Mößinger, K., Brüning, T., Lamanna, F., Belles, M.R., Conrad, C., Kondova, I., et al. (2023) The molecular evolution of spermatogenesis across mammals. Nature 613, 308–316.

Muschick, M., Indermaur, A. & Salzburger, W. (2012) Convergent evolution within an adaptive radiation of cichlid fishes. Current biology 22, 2362–2368.

Ng, A.N.Y., de Jong-Curtain, T.A., Mawdsley, D.J., White, S.J., Shin, J., Appel, B., Dong, P.D.S., Stainier, D.Y.R. & Heath, J.K. (2005) Formation of the digestive system in zebrafish: III. Intestinal epithelium morphogenesis. Developmental biology 286, 114–135.

Ocampo, M., Pincheira-Donoso, D., Sayol, F. & Rios, R.S. (2022) Evolutionary transitions in diet influence the exceptional diversification of a lizard adaptive radiation. BMC ecology and evolution 22, 74.

Orme, D. (2023) The caper package: comparative analysis of phylogenetics and evolution in R.

Papakostas, S., Vøllestad, L.A., Bruneaux, M., Aykanat, T., Vanoverbeke, J., Ning, M., Primmer, C.R. & Leder, E.H. (2014) Gene pleiotropy constrains gene expression changes in fish adapted to different thermal conditions. Nature communications 5, 4071.

Park, J., Levic, D.S., Sumigray, K.D., Bagwell, J., Eroglu, O., Block, C.L., Eroglu, C., Barry, R., Lickwar, C.R., Rawls, J.F., Watts, S.A., Lechler, T. & Bagnat, M. (2019) Lysosome-rich enterocytes mediate protein absorption in the vertebrate gut. Developmental cell 51, 7–20.e6.

Pelster, B., Wood, C.M., Speers-Roesch, B., Driedzic, W.R., Almeida-Val, V. & Val, A. (2015) Gut transport characteristics in herbivorous and carnivorous serrasalmid fish from ion-poor Rio Negro water. Journal of comparative physiology. B, Biochemical, systemic, and environmental physiology 185, 225–241.

Peterson, L.W. & Artis, D. (2014) Intestinal epithelial cells: regulators of barrier function and immune homeostasis. Nature reviews. Immunology 14, 141–153.

Phipson, B., Sim, C.B., Porrello, E.R., Hewitt, A.W., Powell, J. & Oshlack, A. (2022) propeller: testing for differences in cell type proportions in single cell data. Bioinformatics 38, 4720–4726.

Plummer, M., Best, N., Cowles, K. & Vines, K. (2006) CODA: Convergence Diagnosis and Output Analysis for MCMC. R News 6, 7–11.

Pough, F.H., Janis, C.M. & Heiser, J.B. (2013) Vertebrate Life, 9th edition. Pearson, Upper Saddle River, NJ, USA.

Putri, G.H., Anders, S., Pyl, P.T., Pimanda, J.E. & Zanini, F. (2022) Analysing high-throughput sequencing data in Python with HTSeq 2.0. Bioinformatics 38, 2943–2945.

Rajkov, J., El Taher, A., Böhne, A., Salzburger, W. & Egger, B. (2021) Gene expression remodelling and immune response during adaptive divergence in an African cichlid fish. Molecular ecology 30, 274–296.

Revell, L.J. (2024) phytools 2.0: an updated R ecosystem for phylogenetic comparative methods (and other things). PeerJ 12, e16505.

Ricci, V., Ronco, F., Boileau, N. & Salzburger, W. (2023) Visual opsin gene expression evolution in the adaptive radiation of cichlid fishes of Lake Tanganyika. Science advances 9, eadg6568.

Rickelton, K., Zintel, T.M., Pizzollo, J., Miller, E., Ely, J.J., Raghanti, M.A., Hopkins, W.D., Hof, P.R., Sherwood, C.C., Bauernfeind, A.L. & Babbitt, C.C. (2024) Tempo and mode of gene expression evolution in the brain across primates. eLife 13, e70276.

Ritchie, M.E., Phipson, B., Wu, D., Hu, Y., Law, C.W., Shi, W. & Smyth, G.K. (2015) limma powers differential expression analyses for RNA-sequencing and microarray studies. Nucleic acids research 43, e47.

Rivera, C.A., Randrian, V., Richer, W., Gerber-Ferder, Y., Delgado, M.-G., Chikina, A.S., Frede, A., Sorini, C., Maurin, M., Kammoun-Chaari, H., Parigi, S.M., Goudot, C., Cabeza-Cabrerizo, M., Baulande, S., Lameiras, S., et al. (2022) Epithelial colonization by gut dendritic cells promotes their functional diversification. Immunity 55, 129–144.e8.

Rombout, J.H.W.M., Abelli, L., Picchietti, S., Scapigliati, G. & Kiron, V. (2011) Teleost intestinal immunology. Fish & shellfish immunology 31, 616–626.

Ronco, F., Büscher, H.H., Indermaur, A. & Salzburger, W. (2020) The taxonomic diversity of the cichlid fish fauna of ancient Lake Tanganyika, East Africa. Journal of Great Lakes research 46, 1067–1078.

Ronco, F., Matschiner, M., Böhne, A., Boila, A., Büscher, H.H., El Taher, A., Indermaur, A., Malinsky, M., Ricci, V., Kahmen, A., Jentoft, S. & Salzburger, W. (2021) Drivers and dynamics of a massive adaptive radiation in cichlid fishes. Nature 589, 76–81.

Ronco, F. & Salzburger, W. (2021) Tracing evolutionary decoupling of oral and pharyngeal jaws in cichlid fishes. Evolution letters 5, 625–635.

Roux, J. & Robinson-Rechavi, M. (2008) Developmental constraints on vertebrate genome evolution. PLoS genetics 4, e1000311.

RÜber, L. & Adams, D.C. (2001) Evolutionary convergence of body shape and trophic morphology in cichlids from Lake Tanganyika. Journal of evolutionary biology 14, 325–332.

Salzburger, W. (2018) Understanding explosive diversification through cichlid fish genomics. Nature reviews. Genetics 19, 705–717.

Saqui-Salces, M., Huang, Z., Vila, M.F., Li, J., Mielke, J.A., Urriola, P.E. & Shurson, G.C. (2017) Modulation of intestinal cell differentiation in growing pigs is dependent on the fiber source in the diet. Journal of animal science 95, 1179–1190.

Satija, R., Farrell, J.A., Gennert, D., Schier, A.F. & Regev, A. (2015) Spatial reconstruction of single-cell gene expression data. Nature biotechnology 33, 495–502.

Schluter, D. (2000) The Ecology Of Adaptive Radiation. Oxford University Press, London, England.

Sender, R., Weiss, Y., Navon, Y., Milo, I., Azulay, N., Keren, L., Fuchs, S., Ben-Zvi, D., Noor, E. & Milo, R. (2023) The total mass, number, and distribution of immune cells in the human body. Proceedings of the National Academy of Sciences of the United States of America 120, e2308511120.

Shay, T., Jojic, V., Zuk, O., Rothamel, K., Puyraimond-Zemmour, D., Feng, T., Wakamatsu, E., Benoist, C., Koller, D., Regev, A. & ImmGen Consortium (2013) Conservation and divergence in the transcriptional programs of the human and mouse immune systems. Proceedings of the National Academy of Sciences of the United States of America 110, 2946–2951.

Simpson, G.G. (1953) The Major Features Of Evolution. Columbia University Press, New York Chichester, West Sussex.

Singh, P., Irisarri, I., Torres-Dowdall, J., Thallinger, G.G., Svardal, H., Lemmon, E.M., Lemmon, A.R., Koblmüller, S., Meyer, A. & Sturmbauer, C. (2022) Phylogenomics of trophically diverse cichlids disentangles processes driving adaptive radiation and repeated trophic transitions. Ecology and evolution 12, e9077.

Skinnider, M.A., Squair, J.W., Kathe, C., Anderson, M.A., Gautier, M., Matson, K.J.E., Milano, M., Hutson, T.H., Barraud, Q., Phillips, A.A., Foster, L.J., La Manno, G., Levine, A.J. & Courtine, G. (2021) Cell type prioritization in single-cell data. Nature biotechnology 39, 30–34.

Starck, J.M. & Wang, T. (2005) Physiological And Ecological Adaptations To Feeding In Vertebrates. NH: Science Publishers.

Stevens, C.E. (1990) Comparative Physiology Of Vertebrate Digestive Systems. Cambridge University Press, Cambridge, England.

Stuart, T., Butler, A., Hoffman, P., Hafemeister, C., Papalexi, E., Mauck, W.M., 3rd, Hao, Y., Stoeckius, M., Smibert, P. & Satija, R. (2019) Comprehensive integration of single-cell data. Cell 177, 1888–1902.

Subramanian, A., Alperovich, M., Yang, Y. & Li, B. (2022) Biology-inspired data-driven quality control for scientific discovery in single-cell transcriptomics. Genome biology 23, 267.

Sun, Y., Li, H., Liu, Y., Mattson, M.P., Rao, M.S. & Zhan, M. (2008) Evolutionarily conserved transcriptional co-expression guiding embryonic stem cell differentiation. PloS one 3, e3406.

Supek, F., Bošnjak, M., Škunca, N. & Šmuc, T. (2011) REVIGO summarizes and visualizes long lists of gene ontology terms. PloS one 6, e21800.

Szklarczyk, D., Kirsch, R., Koutrouli, M., Nastou, K., Mehryary, F., Hachilif, R., Gable, A.L., Fang, T., Doncheva, N.T., Pyysalo, S., Bork, P., Jensen, L.J. & von Mering, C. (2022) The STRING database in 2023: protein– protein association networks and functional enrichment analyses for any sequenced genome of interest. Nucleic acids research 51, D638–D646.

Takeyama, T. (2021) Feeding ecology of Lake Tanganyika cichlids. In The Behavior, Ecology and Evolution of Cichlid Fishes (eds M.E. Abate & D.L.G. Noakes), pp. 715–751.

Tanay, A. & Sebé-Pedrós, A. (2021) Evolutionary cell type mapping with single-cell genomics. Trends in genetics: TIG 37, 919–932.

Tibbetts, I.R. (1997) The distribution and function of mucous cells and their secretions in the alimentary tract of *Arrhamphus sclerolepis krefftii*. Journal of fish biology 50, 809–820.

Tosches, M.A., Yamawaki, T.M., Naumann, R.K., Jacobi, A.A., Tushev, G. & Laurent, G. (2018) Evolution of pallium, hippocampus, and cortical cell types revealed by single-cell transcriptomics in reptiles. Science 360, 881–888.

Tóth, B., Ben-Moshe, S., Gavish, A., Barkai, N. & Itzkovitz, S. (2017) Early commitment and robust differentiation in colonic crypts. Molecular systems biology 13, 902. EMBO.

Ullrich, K.K. & Glytnasi, N.E. (2023) oggmap: a Python package to extract gene ages per orthogroup and link them with single-cell RNA data. Bioinformatics 39, btad657.

Venditti, C., Meade, A. & Pagel, M. (2011) Multiple routes to mammalian diversity. Nature 479, 393–396.

Wagner, C.E., McIntyre, P.B., Buels, K.S., Gilbert, D.M. & Michel, E. (2009) Diet predicts intestine length in Lake Tanganyika’s cichlid fishes. Functional ecology 23, 1122–1131.

Wang, B., Starr, A.L. & Fraser, H.B. (2024) Cell type-specific cis-regulatory divergence in gene expression and chromatin accessibility revealed by human-chimpanzee hybrid cells. eLife 12, RP89594.

Wittkopp, P.J. & Kalay, G. (2011) Cis-regulatory elements: molecular mechanisms and evolutionary processes underlying divergence. Nature reviews. Genetics 13, 59–69.

Xia, J.H., Li, H.L., Li, B.J., Gu, X.H. & Lin, H.R. (2018) Acute hypoxia stress induced abundant differential expression genes and alternative splicing events in heart of tilapia. Gene 639, 52–61.

Xu, C., Li, E., Suo, Y., Su, Y., Lu, M., Zhao, Q., Qin, J.G. & Chen, L. (2018) Histological and transcriptomic responses of two immune organs, the spleen and head kidney, in Nile tilapia (*Oreochromis niloticus*) to long-term hypersaline stress. Fish & shellfish immunology 76, 48–57.

Young, M.D. & Behjati, S. (2020) SoupX removes ambient RNA contamination from droplet-based single-cell RNA sequencing data. GigaScience 9, giaa151.

Zhang, J. (2023) Patterns and evolutionary consequences of pleiotropy. Annual review of ecology, evolution, and systematics 54, 1–19.

Zihler, F. (1981) Gross morphology and configuration of digestive tracts of Cichlidae (teleostei, Perciformes): Phylogenetic and functional significance. Netherlands journal of zoology 32, 544–571.

Zwick, R.K., Kasparek, P., Palikuqi, B., Viragova, S., Weichselbaum, L., McGinnis, C.S., McKinley, K.L., Rathnayake, A., Vaka, D., Nguyen, V., Trentesaux, C., Reyes, E., Gupta, A.R., Gartner, Z.J., Locksley, R.M., et al. (2024) Epithelial zonation along the mouse and human small intestine defines five discrete metabolic domains. Nature cell biology 26, 250–262.

